# Decoding cnidarian cell type gene regulation

**DOI:** 10.1101/2025.07.01.662323

**Authors:** Anamaria Elek, Marta Iglesias, Lukas Mahieu, Grygoriy Zolotarov, Xavier Grau-Bové, Stein Aerts, Arnau Sebé-Pedrós

## Abstract

Animal cell types are defined by differential access to genomic information, a process orchestrated by the combinatorial activity of transcription factors that bind to *cis*-regulatory elements (CREs) to control gene expression. However, the regulatory logic and specific gene networks that define cell identities remain poorly resolved across the animal tree of life. As early-branching metazoans, cnidarians can offer insights into the early evolution of cell type-specific genome regulation. Here, we profiled chromatin accessibility in 60,000 cells from whole adults and gastrula-stage embryos of the sea anemone *Nematostella vectensis.* We identified 112,728 CREs and quantified their activity across cell types, revealing pervasive combinatorial enhancer usage and distinct promoter architectures. To decode the underlying regulatory grammar, we trained sequence-based models predicting CRE accessibility and used these models to infer ontogenetic relationships among cell types. By integrating sequence motifs, transcription factor expression, and CRE accessibility, we systematically reconstructed the gene regulatory networks that define cnidarian cell types. Our results reveal the regulatory complexity underlying cell differentiation in a morphologically simple animal and highlight conserved principles in animal gene regulation. This work provides a foundation for comparative regulatory genomics to understand the evolutionary emergence of animal cell type diversity.

## Introduction

In multicellular animals, cell type-specific gene expression is orchestrated by transcription factors (TFs), which recognize specific sequence motifs located within *cis*-regulatory elements (CREs) such as gene promoters and enhancers. These TF-CRE networks ultimately interpret genomic information in each cell, determining the transcriptional state of individual genes and collectively shaping specific gene regulatory networks (GRNs). By measuring the transcriptional output of these gene programs, single-cell transcriptomics provided unprecedented insights into the molecular diversity of cell types across animal lineages^1–9^. However, our understanding of the structure and logic of the regulatory programs that define cell types remains limited for most species, except for fruit fly^10–12^ and vertebrates^13–17^. The recent development of single-cell chromatin accessibility assays (scATAC-seq)^18^, together with the generation of high-quality genomes and gene expression data, has opened new opportunities to study whole-organism, cell type-specific gene programs in non-model species.

Cnidarians (anemones, corals, and jellyfish) can offer insights into the evolution of animal regulatory complexity, given their key phylogenetic position as sister group to all bilaterian animals. Notably, cnidarian genomes show hallmarks of bilaterian gene regulation such as distal enhancer elements^19^. Furthermore, while historically considered simple animals with relatively few cell types, single-cell transcriptomics studies have revealed that cnidarians encode a diverse repertoire of cell types^2,7,20–23^. To understand the genomic basis of this cell diversity, we systematically dissected cell type *cis*-regulatory programs in the sea anemone *Nematostella vectensis* (**Fig. 1a)**, including cell-specific CRE landscapes, regulatory sequences, and GRNs.

**Figure 1.**
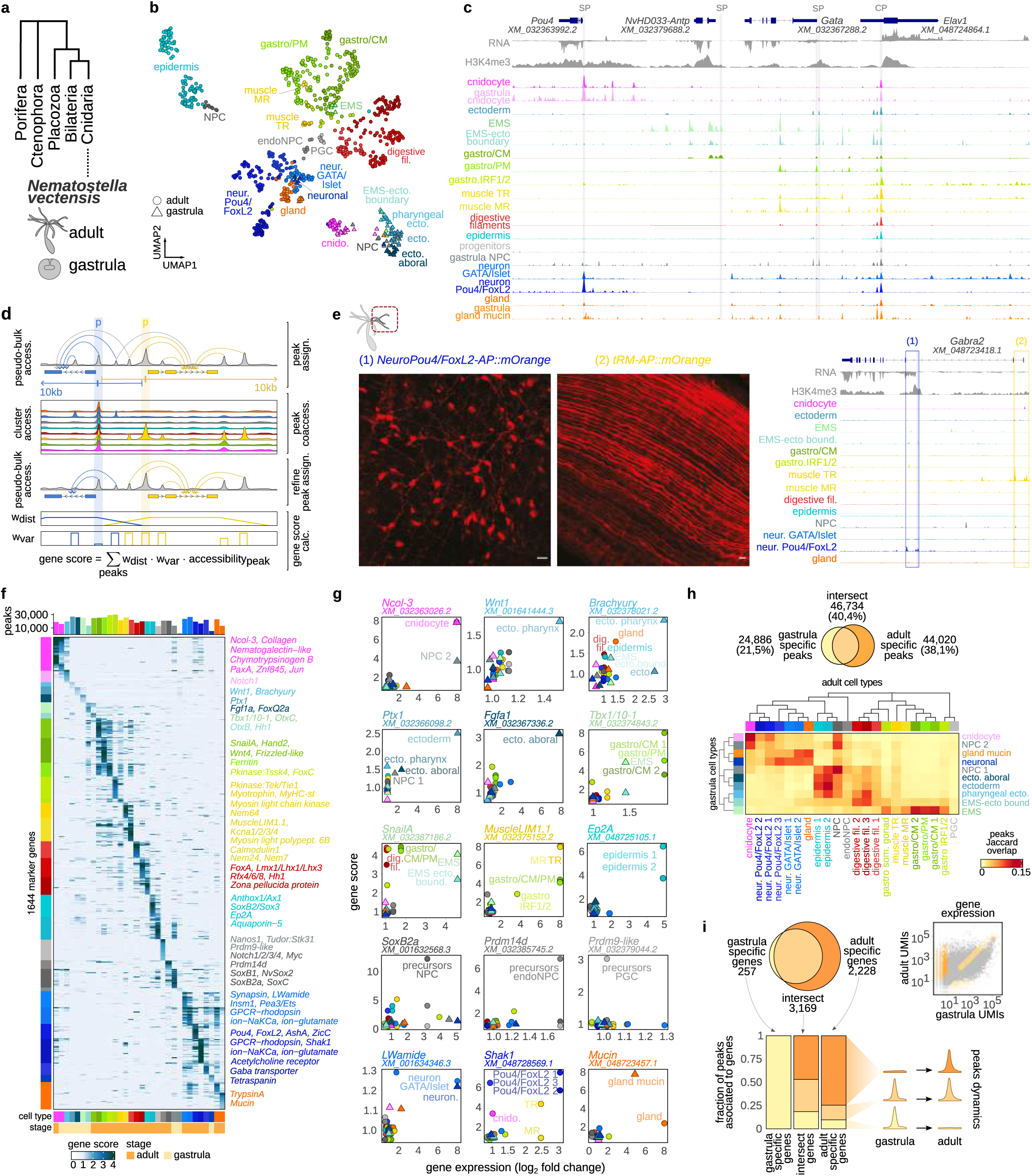
Cell type-specific chromatin landscapes in *Nematostella vectensis.* **a,** *Nematostella* phylogenetic position. **b,** UMAP 2D projection of scATAC metacells, colored by cell type, with broad cell type labels. Gastrodermis/parietal muscle (gastro/PM), gastrodermis/circular muscle (gastro/CM), mesentery retractor muscle (MR); tentacle retractor muscle (TR), ectoderm (ecto.), and endomesoderm (EMS). **c,** Example regulatory landscapes for selected genes. Forward and reverse RNA signal is shown above and below baseline, respectively. Promoter peaks are highlighted with grey boxes. SP specific promoter, CP constitutive promoter. **d,** Peak assignment and gene accessibility score calculation strategy. Peaks up to 10 kb are assigned to genes, unless they are downstream of another gene’s promoter. When a peak is assigned to more than one gene, peak-peak co-accessibility is used to refine peak assignment. Gene score is then calculated as a sum of the accessibility of peaks assigned to a gene, weighted by distance from the TSS (w_var_) and peak variability across clusters (w_var_). **e,** Transgenic reporter validation of *Gabra2* alternative promoters. Images correspond to the tentacle region showing reporter expression in neurons and longitudinal muscle fibers (left) corresponding to the regulatory regions highlighted in the genome browser (right). **f,** Heatmap of gene scores for marker genes across cell types. Color code for genes indicates the cell type where gene has the highest score. Selected known markers are highlighted on the right. **g,** Comparison between gene accessibility scores and gene expression levels for selected marker genes. **h,** Euler diagram showing the total number of overlapping peaks (accessibility FC > 1.5) between the two life stages (top) and heatmap representing peak overlap between adult and gastrula cell types (bottom). Rows and columns are clustered based on peak overlap between cell types within each life stage. **i,** CRE dynamics across development. We first select genes expressed in adult and/or gastrula (Euler diagram, top) and then analyze the accessibility dynamics of CREs associated to these three gene groups across development (bottom).

## Results

### *Nematostella* cell type-specific chromatin landscapes

To define CRE usage across *Nematostella* cell types, we profiled chromatin accessibility in 51,866 adult and 6,882 gastrula-stage single-cells using 10x Genomics scATAC-seq (**Fig. 1b, Extended Data** Fig. 1a**-c**). We sequenced libraries to an average of 2,606 reads/cell and obtained a median of 908 fragments/cell (873 in adult, 1,362 in gastrula). Cells were grouped based on their accessibility profiles into metacells^24^, which served as the basic units for downstream analyses (**Fig. 1b, Extended Data** Fig. 1d, e). This resulted in 693 metacells for the adult stage and 95 for the gastrula stage, with each metacell containing a median of 67 and 53 single cells, respectively. We then clustered metacells using neighbor joining based on their accessibility profiles (**Extended Data** Fig. 1f-h). We identified chromatin accessibility peaks using cluster-level aggregated pseudobulk ATAC-seq signal and iteratively merged overlapping peaks^25^, generating a catalogue of 112,728 CREs across the 269Mb *Nematostella* genome (**Fig. 1c and Extended Data** Fig. 3d). We assigned peaks to genes based on their distance to transcription start sites (TSS) and co-variation across cell types (**Fig. 1d**) and we annotated scATAC-seq cell clusters using previously defined scRNA-seq cell types^2,26^ (**Extended Data** Fig. 1i-k). To achieve this, we calculated an ensemble gene accessibility score as a weighted sum of peak accessibility for each gene (**Fig. 1d**) and correlated these scores with gene expression to match scATAC-seq clusters to scRNA-defined cell types. This analysis resulted in 32 annotated cell clusters (22 in the adult, 10 in the gastrula), each with both specific and combinatorial gene accessibility patterns (**Fig. 1b**) and with cluster-specific accessible CREs ranging from 2,000 to 30,000 (median 21,156 CREs). To validate these cell type-specific CREs, we generated transgenic reporter lines for two predicted alternative promoters of the *Gabra2* gene. The two promoters drove expression in either the tentacle retractor muscle or tentacle neurons, recapitulating their respective accessibility profiles (**Fig. 1e**).

Adult cell clusters included eight previously described broad adult cell types^2^, characterized by high CRE accessibility around known markers (**Fig. 1f, g, Extended Data** Fig. 2) such as *Ncol-3* (cnidocytes), *MuscleLIM* protein (retractor muscle), *EP2A* (epidermis), and *Shak3* ion channel (Pou4/FoxL2 neurons). We also identified three distinct clusters of adult progenitor cells. One represents adult neurosecretory progenitors (NPCs) characterized by differential accessibility near TF genes like *SoxC*, *SoxB2a,* and *Ath-like*^26,27^. Another, that we termed endodermal NPCs (endo-NPCs), exhibited accessibility near *Prdm14d*, a marker for endodermal neurogenesis^28^. The third precursor cluster likely represented primordial germ cells (PGCs), characterized by the differential accessibility near *Prdm9*^29^. Gastrula-stage cell clusters included both differentiated cell types, such as gland cells, cnidocytes, and neurons, as well as progenitor cells, such as NPCs. We also identified the main germ layers and spatial territories within the gastrula ^26,30^: ectoderm and aboral ectoderm, showing *Ptx1*^26^ and *Ffg1a*^31^ accessibility; endomesoderm (EMS), showing *Tbx1/10-1*^30^ and *SnailA*^30^ accessibility; and pharyngeal ectoderm, showing *Brachyury*^30^, *FoxA* and *Wnt1*^32^ differential accessibility (**Fig. 1g, Extended Data** Fig. 2).

We then compared CRE usage between adult and gastrula cell types, identifying 46,734 shared CRE (40,4%) between adult and gastrula (**Fig. 1h**). Comparisons of CRE accessibility revealed strong similarities between neurons, cnidocytes, and gland cells, as well as between NPCs at both stages (**Fig. 1h**). Additionally, the CRE landscapes of gastrula germ layers showed resemblances to some adult cell types, including similarities between EMS and gastrodermal/mesenteric retractor muscles, between pharyngeal ectoderm and digestive filaments, and between ectoderm and epidermis. While the precise developmental trajectories of these tissues remain to be fully characterized, these patterns align with the hypothesis of three germ layers in *Nematostella* as proposed by Steinmetz et al.^30^. Finally, we found that approximately 25% of the genes exclusively expressed in adult *Nematostella* had CREs accessible already at the gastrula stage, including 10% of CREs only accessible in gastrula stage (**Fig. 1i**). This is consistent with findings in other species, where chromatin accessibility often precedes transcriptional activation^33^, and could reflect the early activity of pioneer TFs (**Fig. 1i**). Overall, our *Nematostella* single-cell accessibility atlas represents a comprehensive inventory of cell type-specific *cis*-regulatory landscapes in a non-bilaterian animal. This atlas is available for exploration through an interactive database and genome browser: https://sebelab.crg.eu/nematostella-cis-regulatory-atlas/ and https://sebelab.crg.eu/nematostella-cis-reg-jb2

### Cnidarian gene regulatory architecture

We next investigated the different CRE configurations associated with *Nematostella* genes. First, we classified CREs into promoters and non-promoters (which we termed *enhancers*) using a combination of distance to TSS, histone post-translational modifications (H3K4me3), and 5ʹ scRNA-seq data (**Extended Data** Fig. 3a). Among the 58,954 CREs identified in adult cell types, we classified 21,344 (36%) as promoters and 37,610 (64%) as enhancers (**Fig. 2a, Extended Data** Fig. 3b). These proportions are similar to those observed in *Drosophila*, which has 27% promoters and 73% enhancers, whereas in mice, the fraction of promoters among scATAC-defined CREs is significantly smaller (5% vs. 95% enhancers). In *Nematostella*, enhancers are predominantly located in intergenic regions (38.8%), followed by intronic regions (26.8%). Enhancers in mouse also tend to be found in intergenic regions (46%), while in *Drosophila* 49% are intronic and 22% are intergenic (**Fig. 2a**).

**Figure 2.**
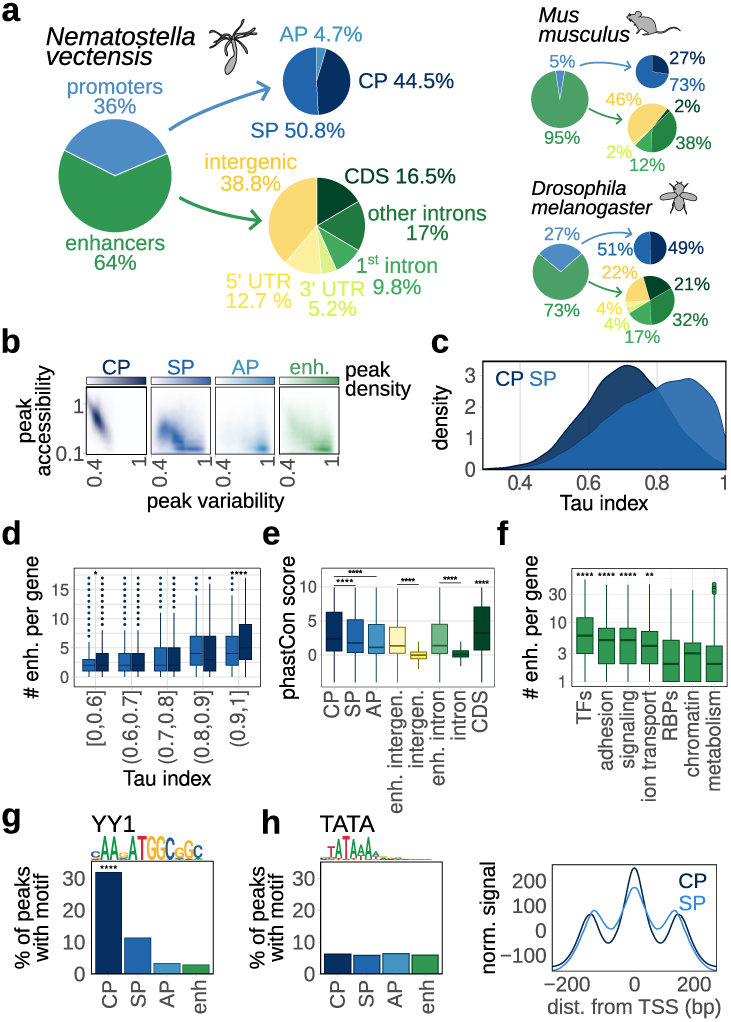
Pre-bilaterian gene regulatory architecture. **a,** Fraction of CREs classified as promoters and enhancers. Promoters are further classified as constitutive promoters (CP), specific promoters (SP) and alternative promoters (AP). Enhancers are classified based on their overlap with different genomic regions. The same is shown for *Nematostella* (top), mouse (bottom left) and *Drosophila melanogaster* (bottom right). **b,** Comparison of accessibility *versus* variability across cell clusters for different CRE classes. **c,** Cell type expression specificity (Tau index) for genes with CP and SP. **d,** Number of enhancers for group of genes with CP or SP and different levels of expression specificity (Tau index bins). **e,** Sequence conservation (phastCon score) of different promoter classes and enhancers overlapping intergenic and intronic regions. Conservation of intergenic and intronic regions not overlapping predicted enhancers is shown for comparison. **f,** Number of enhancers for different functional gene sets. RNA-binding proteins (RBPs). **g,** Fraction of peaks with YY1 motif in different CRE classes. **h,** Fraction of peaks with TATA motif in different CRE classes (left) and aggregated ATAC signal y around promoters (right). * p < 0.05, **** p < 0.0001 (Wilcoxon test).

Focusing on promoters, we identified approximately half (44.5%) as constitutively accessible across all cell types (Constitutive Promoters, CP), while roughly another half (50.8%) were cell type-specific (Specific Promoters, SP). A smaller fraction (4.7%) represented alternative promoters (AP) of the same gene accessible in different cell types (**Fig. 2a and Extended Data** Fig. 3a). These proportions are similar in *Drosophila* (51% SP vs. 49% CP), whereas in mouse SP promoters are more frequent (73% SP vs 27% CP). *Nematostella* CPs are, on average, showed higher and less variable accessibility compared to SP, AP and enhancers (**Fig. 2b**) — a pattern similar to what has been observed in *Drosophila*^33^. Additionally, CPs were generally associated with genes expressed across multiple cell types, whereas genes with SPs tended to exhibit more restricted, cell type-specific expression (**Fig. 2c**). Regardless of promoter type, genes with cell type-specific expression were linked to increased number of associated enhancer elements (**Fig. 2d**). Comparing CRE sequence conservation across cnidarian genomes, we found that CPs are significantly more evolutionarily conserved than SPs, APs, or enhancers (**Fig. 2e**). Furthermore, TFs represent the gene class with the highest number of associated enhancers (**Fig. 2f**).

We also examined sequence motifs enriched in different promoter types and found that YY1 motif was strongly enriched in CP (**Fig. 2g**). *YY1* is a metazoan-specific TF that has been involved in enhancer-promoter contacts in different cell types^34^, suggesting that *Nematostella* CPs may rely on this factor for integrating regulatory signals from their associated enhancers. In bilaterian animals, adult cell type-specific promoters —often called Type I promoters^35,36^— are characterized by the presence of TATA motifs and fuzzy nucleosomes. In contrast, *Nematostella* SP have well-positioned flanking nucleosomes and lacked TATA motifs (**Fig. 2h, Extended Data** Fig. 3f), suggesting that this class of promoters may be a bilaterian-specific feature. These findings offer a comprehensive perspective on the landscape of cell type-specific gene regulation in a non-bilaterian animal. Our results highlight similarities to bilaterians, such as CRE type proportions and genomic distributions, while also revealing key differences, including the absence of TATA-containing Type I promoters.

### *Nematostella cis*-regulatory programs

Having defined the CRE accessible in different cell types, we next sought to identify the key TFs and *cis*-regulatory sequences in each cell type. To determine which sequence motifs are important for CRE accessibility, we employed two complementary approaches: (1) calculating motif enrichments in accessible CREs using both *de novo* discovered and known motif collections (**Extended Data** Fig. 4a), and (2) training sequence-to-function machine learning models that explain the relationship between sequence features and accessibility^37–40^, to then extract important model features and discovering motifs^41,42^ (**Extended Data** Fig. 5). To reduce redundancy in motif annotations, we grouped similar motifs into broader archetypes^43^, and then compared the motif collections obtained with each method (**Extended Data** Fig. 6a-i). Motif enrichment analyses uncovered a larger number of motifs (1,292) compared to sequence models (637). This discrepancy could be explained because sequence models prioritize motifs that are predictive of accessibility patterns rather than capturing an exhaustive set of all enriched motifs. However, up to 30% of motifs identified by sequence models were absent from enrichment analyses, suggesting that sequence models offer higher sensitivity and can detect important motifs with fewer genome-wide binding sites.

Beyond motif discovery, we also leveraged sequence models to investigate cell type-specific CRE codes, considering both motif composition (lexicons) and the combinatorial rules governing motif arrangement, orientation and spacing (syntax). For example, in adult cnidocytes, the most common motif grammar contained Pou4 in combination with E-box bHLH, Fox, and zf-C2H2 motifs (**Fig. 3a, Extended Data** Fig. 6j). Across all cell types, we identified 15–36 key motifs per cell type, and each CRE containing a median of 3–4 motif instances (**Fig. 3b**). The co-occurrence of TF binding motifs within CREs ranged from 10% to 75% depending on the cell type (**Fig. 3c**). When analyzing motif combinations, we found that most motif pairs and triplets exhibited flexible order and orientation (**Fig. 3d**), with only a few exceptions involving YY1 and zf-C2H2 binding sites. This pattern observed in *Nematostella* is compatible with a billboard-like model of TF binding sites^44^.

**Figure 3.**
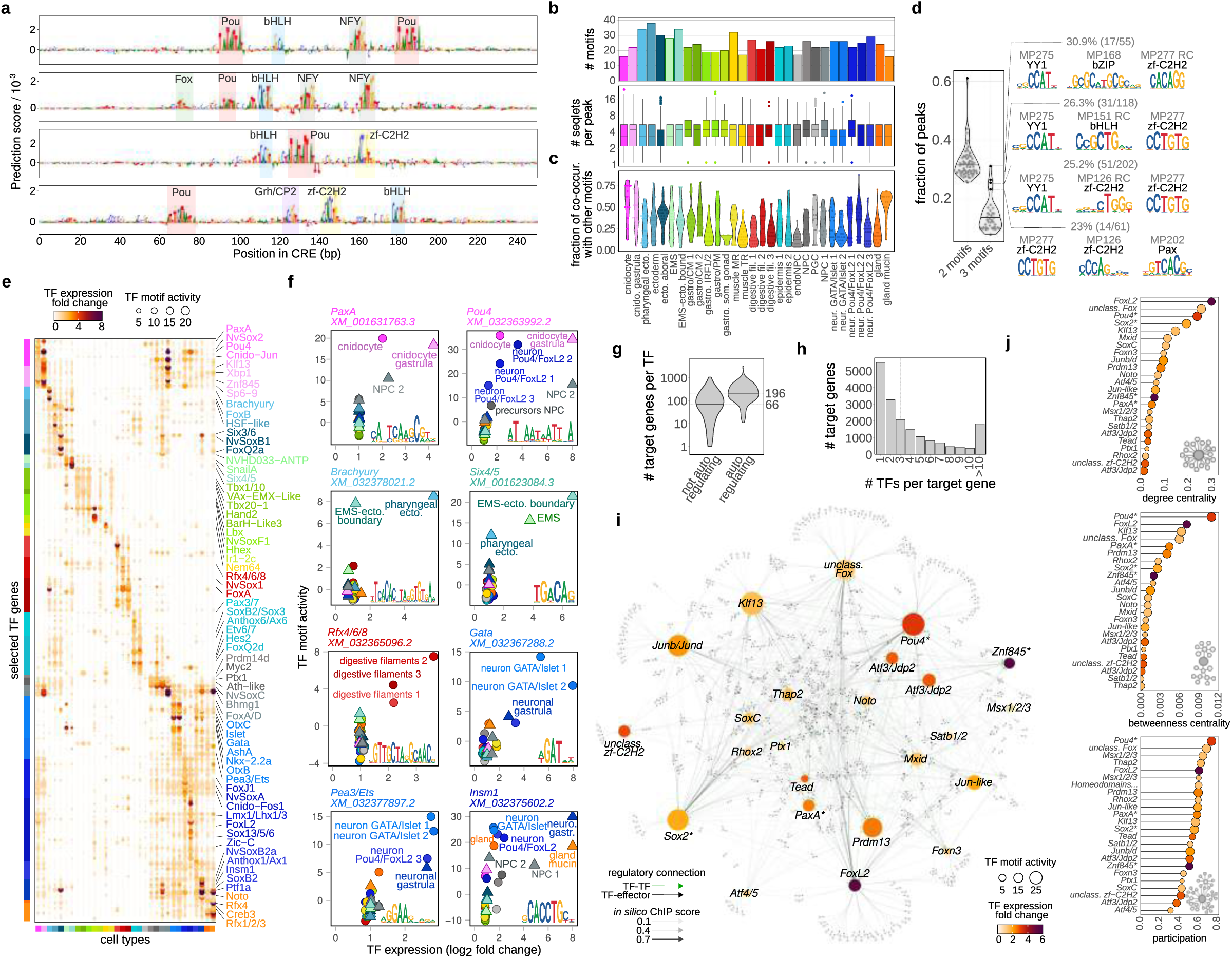
*Nematostella* cell type regulatory programs. **a,** Nucleotide importance scores for four representative cnidocyte CREs, highlighting detected TF motifs. **b,** Total number of motifs learned from sequence models per cell type (top), and total number of motif instances (seqlets) in each peak per cell type (bottom). **c,** Fraction of TF motif co-occurrences per cell type. We define two motifs as co-occurring when they appear non-overlapping in the same CRE – e.g., co-occurrence fraction of 0.5 would mean that 50% of CREs where TF motif is identified also have at least one more motif of another TF. **d,** Frequency distribution of motif pairs (doublets) and triplets that co-occur in more than 50 CREs. The violin plot (left) shows the fraction of peaks (out of all peaks with given motif combination) in which motifs appear in specific order and orientation. The examples of motif triplets that deviate from flexible ordering and orientation are highlighted (right). **e,** Dotmap showing TF motif activity (dot size) and expression (colorscale) for selected variable TFs across cell types. Cell types are color coded as in Fig.1. Color code for TFs indicates the cell type where a TF has the highest motif activity. **f,** Examples of correlated TF expression and TF motif activity. **g,** Number of target genes per TF, shown separately for (predicted) self-regulating TFs (median=196) and not-self regulating TFs (median=66). **h,** Number of TF regulators per target gene (median=3). **i,** Inferred GRN for cnidocytes in adult *Nematostella*. TF nodes are colored by expression, scaled by TF motif activity and labeled; target genes are indicated by small grey dots. Width and transparency of connections represents interaction strength (*in silico* ChIP binding score). **j,** Network centrality metrics for the top TFs in the inferred cnidocyte GRN.

To further link CRE sequences to TF function, we assigned motifs to specific *Nematostella* TFs using a combination of orthology, sequence similarity-based motif transfer^45^, and correlations between TF expression and motif accessibility (see Methods) (**Extended Data** Fig. 4b-d). This analysis enabled us to predict candidate binding motifs for 96% (571/590) of expressed/accessible TFs in *Nematostella*. Then, we compared TF expression to the aggregated accessibility of the assigned motif in each cell type (TF motif activity)^46^, observing good agreement between TF expression and TF activity (**Fig. 3e**). For example, we found that *PaxA* was specifically expressed and active in cnidocytes, *FoxA* and *Rfx4/6/8* in digestive filaments, *Hes2* in ectodermal cells, and *FoxQ2d* in epidermis. *Pou4* is expressed and active in cnidocytes and one broad type of neurons; while *Gata*, *Islet* and *OtxC* are active in the other broad neuronal type (**Fig. 3f, Extended Data** Fig. 7).

We integrated CRE accessibility with TF motif binding scores and gene expression to infer cell type-specific gene regulatory networks (GRNs). We used *in silico* ChIP^47^ to link TFs to target CREs that both contained a high-scoring motif for the TF and exhibited accessibility patterns correlated with TF expression across cell types. This allowed us to reconstruct a global TF-CRE network, which we then refined for each cell type based on TF motif activity, TF expression, and target peak accessibility. In the global GRN model, TFs lacking self-regulation were predicted to target a median of 68 genes, whereas self-regulating TFs targeted a median of 212 genes (**Fig. 3g**). This suggests that self-regulating TFs may control larger networks of effector genes, contributing to long-term maintenance of cell functions. From a complementary perspective, each effector gene is predicted to be regulated by a median of 3 TFs (**Fig. 3h**). Within a cell type, TFs were found to regulate very different sets of genes (median overlap between predicted targets 3%) (**Extended Data** Fig. 8a). Across cell types, TFs tend to regulate distinct sets of genes as a function of the number of cell types in which these TFs are active (**Extended Data** Fig. 8b). The analysis of GRN structure also highlights important TF for cell type identity (**Fig. 3i**, **Extended Data** Fig. 8d-i). For example, the reconstructed GRN for adult cnidocytes (**Fig. 3j**) indicates that *FoxL2*^2^ is the TF with most regulatory connections (i.e. highest degree of centrality), *Pou4*^48^ is the TF bridging most submodulies in the network (i.e. highest betweenness centrality) and *NvSox2*^49^ is the global regulator with connections spread most evently across different submodules (i.e. highest participation). In each cell type, we also identified a subset of TFs with predicted self-regulation, for example *FoxL2*, *Pou4* and *NvSox2* in the case of cnidocytes (**Fig. 3j**).

### Cell type relationships defined by regulatory characters

We finally explored the relationships between the identified *Nematostella* cell clusters by comparing different regulatory features. We first grouped cell clusters based on Euclidean distances between gene accessibility profiles (gene scores; **Fig. 4a**), which we expected to largely reflect effector gene activity in a manner analogous to gene expression. This analysis revealed that functionally related cell types tended to cluster together, for example adult muscle cell types (fast contracting retractor muscles, and slow contracting parietal and circular muscles), as well as a group composed of neurosecretory cells (cnidocytes, neurons, and gland/secretory cells) alongside epidermal cells and neuronal progenitors. In contrast, clustering based on the overlap of accessible CREs (**Fig. 4b)** in resulted in a different grouping: tentacle retractor muscle (TR) cells clustered with epidermal cells and adult neuronal progenitor cells (NPCs), while the remaining muscle cell types grouped together with gastrula endomesodermal (EMS) cells. A similar pattern, consistent with known ontogenetic relationships, was observed when we compared cells based on *cis*-regulatory sequence similarity, using AUC values derived from gkmSVM classifiers performance across cell types (**Fig. 4c)**. This analysis revealed the strongest cross-stage associations. For instance, gastrula ectodermal cell types clustered with known ectodermally derived adult cell types such as epidermis, NPCs, cnidocytes^50^ and TR muscle^51^, along with Pou4/FoxL2-expressing neurons. Separately, gastrula EMS cells clustered with endomesodermally derived adult muscle types—including mesenteric retractor, circular, and parietal muscles^51^. Another cross-stage association included gastrula pharyngeal ectoderm cells clustering with digestive filaments^52^. Interestingly, adult gland/secretory cells and GATA/Islet positive neurons formed a distinct cluster that was more similar to the group of endomesodermal and pharyngeal derivatives than to Pou4/FoxL2-positive neurons and cnidocytes. This may suggest the existence of developmentally distinct populations of enteric and ectodermal/epidermal neurons in *Nematostella*^53,54^.

**Figure 4.**
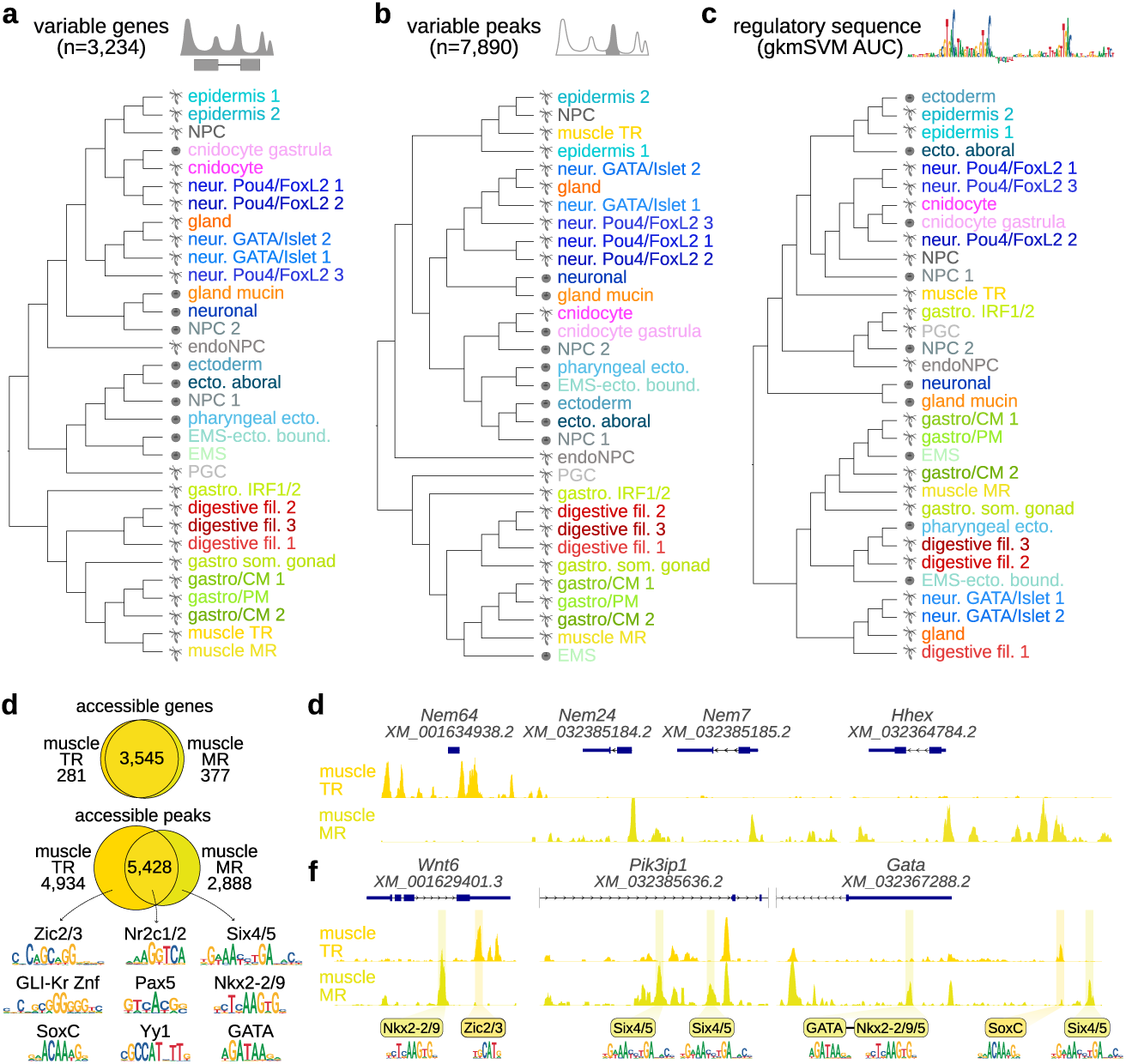
Comparing *Nematostella* cell type regulatory identities. **a,** Neighbor-joining (NJ) cell type tree based on accessibility scores for 3,234 variable genes. **b,** NJ cell type tree based on shared accessibility of 7,980 variable peaks. **c,** NJ Cell type tree based on regulatory sequence similarity, based on AUC values obtained applying cell type gkm-SVM classifiers between cell types. **d,** Euler diagram showing the overlap of genes (top, based on gene scores) and peaks (bottom) accessible in two retractor muscles (MR mesentery retractor, TR tentacle retractor). Top enriched motifs in each group of peaks. **e,** CRE accessibility around key TFs in TR and MR muscle cells. **f,** Examples of genes with shared accessibility but different set of accessible peaks in two retractor muscles. Occurrences of enriched motifs shown in E are shown below the coverage tracks.

The different affinities of the two retractor muscle cell types (tentacle *versus* mesenteric) have implications for the evolution of their expression programs. Distinct tentacle (TR) and mesenteric retractor (MR) muscle cell types were recently identified by Cole et al.^55^ as transcriptionally highly similar cell types but regulated by different bHLH TFs (*Nem64* in TR, and *Nem7* and *Nem24* in MR). Consistent with this finding, TR and MR form clearly separate clusters in our scATAC-seq data (**Fig. 1b**) and and exhibit differential chromatin accessibility around the *Nem64*, *Nem7* and *Nem24* loci. (**Fig. 4e**). TR has been proposed to arise from ectodermal precursors through co-option of the mesenteric MR program—a process thought to be mediated by the emergence of the *Nem64* paralog^55^. To investigate this hypothesis, we compared TR-MR similarity at two levels: genes and CREs. As expected, given their high transcriptional similarity, gene scores for TR and MR were largely overlapping, indicating that many active effector genes are shared between the two (**Fig. 4d**). In contrast, there was far less overlap at the level of individual CREs (**Fig. 4d**). These non-overlapping peaks contain distinct sets of TF motifs, which helps explain the TR-MR distinctiveness at the level of *cis*-regulatory sequences (**Fig. 4c**). These results indicate that albeit most active genes in TR and MR muscle are the same, they are regulated through different CREs that can interpret different uptstream regulatory states (**Fig. 4f**). Thus, our data does not fit with the idea of a simple co-option of the MR cell type program via recruitment of one TF paralog (*Nem64*) into an upstream ectodermal GRN. Instead, our results indicate that many shared muscle effector genes are controlled by distinct CREs, likely reflecting their activation in ontogenetically distinct precursors—ectodermal for TR and endomesodermal for MR.

## Discussion

Here, we present a whole-organism single-cell chromatin accessibility atlas for the cnidarian *Nematostella vectensis*. This atlas allowed us to dissect the regulatory logic underlying cell type-specific gene expression in cnidarians. We identified 112,728 CREs across the 269 Mb *Nematostella* genome, including 91,362 putative enhancers (i.e., non-promoter CRE). This number substantially exceeds previous estimates and approaches the number of CREs reported in *Drosophila*, which has a similar genome size (180Mb).

We identified key TFs associated with each cell identity by analyzing their expression, aggregated motif accessibility, and regulatory influence. In parallel, we defined the *cis*-regulatory motif grammars that characterize cell type-specific CREs. By integrating TF activity with CRE accessibility and motif composition, we inferred gene regulatory network (GRN) models for major *Nematostella* cell types, enabling systematic analysis of GRN structure and composition. These analyses reveal the intricate regulatory logic that governs cell type–specific gene programs in a morphologically simple, non-bilaterian animal.

Our findings further show that while effector gene usage groups functionally similar cell types within each developmental stage, regulatory features can reveal ontogenetic relationships between cell types. For instance, GATA/Islet-expressing neurons exhibit regulatory sequence similarities with endomesodermal and pharyngeal derivatives, clearly distinguishing them from the ectodermally-associated Pou4/FoxL2 neurons. This suggests a possible enteric origin for this broad class of neurons in *Nematostella*. This analysis also sheds light into the co-option process underlying ectodermally-derived muscles in *Nematostella* tentacles. Here, the activation of a similar set of fast muscle effector genes occurs through largely distinct CREs and regulatory sequence information, even for the same target genes. This suggests that developmental homoplasy may not result merely from the duplication and redeployment of a terminal selector TF in a different germ layer (in this case, the ectoderm). Instead, such convergent activation of effector programs may require access to distinct regulatory states, such as those mediated by pioneer TFs that establish CRE accessibility.

Our *cis*-regulatory atlas moves beyond conventional transcriptome-based cell type characterization by analyzing regulatory traits that define cell type identities in *Nematostella,* such as CREs sequence motif composition, active TFs, and GRN architecture. We anticipate that applying similar approaches in other organisms will further advance our understanding of animal genome regulation and serve as a powerful tool for resolving cell type evolutionary relationships across species^56^.

## Data and code availability

Raw and processed files will be available in GEO repository under accession number GEO: GSE294388. In addition, the atlas can be explored in interactive database: https://sebelab.crg.eu/nematostella-cis-regulatory-atlas/ and also in an interactive genome browser: https://sebelab.crg.eu/nematostella-cis-reg-jb2 Scripts to reproduce the data processing and downstream analysis are available in GitHub: https://github.com/sebepedroslab/nvec-scatac. Unless otherwise specified, scripts are based on R version 4.2.2 and Python version 3.8.10, and the language-specific libraries specified in the Methods section.

## Acknowledgements

We thank Iana Kim, Alex de Mendoza, Sean Montgomery, and Nacho Maeso for critical comments on the manuscript, as well as all members of the Sebe-Pedros group for discussion and suggestions. We thank Fabian Rentzsch for access to *Nematostella* Elav1::mOrange transgenic line. Research in A.S-P. group was supported by the European Research Council (ERC-StG 851647) and the Spanish Ministry of Science and Innovation (PID2021-124757NB-I00). We also acknowledge support of the Spanish Ministry of Science and Innovation to the EMBL partnership, the Centro de Excelencia Severo Ochoa and the CERCA Programme (Generalitat de Catalunya). A.E. was supported by FPI PhD fellowship from the Spanish Ministry of Science and Innovation. M.I. was supported by the European Union’s H2020 research and innovation program under Marie Skłodowska-Curie INTEPiD co-fund agreement 75442. X.G-B. was supported by the European Union’s H2020 research and innovation program under Marie Skłodowska-Curie grant agreement 101031767.

## Methods

### Nematostella culture

The *Nematostella vectensis* culture is derived from CH2 males and CH6 females^57^. Adult polyps were maintained at 18°C in 1/3 filtered seawater (*Nematostella* medium, NM), and spawned by a temperature and light shock^58^. Fertilized egg packages were treated with a 3% L-cysteine in NM solution to remove the egg jelly. Embryos were raised at 21°C until mid-gastrula stage (26 hours post fertilization, hpf) and collected based on their morphology.

### Sample preparation for single-cell experiments

Depending on the sample input and single-cell omics protocol, different approaches were used to obtain single cell suspensions as described below:

*<u>1) Whole-gastrula scATAC-seq.</u>* Embryos were washed twice in cold PBS before nuclei isolation and permeabilization. Nuclei from 300 pooled gastrula were isolated in 300ml of 1x OmniATAC lysis buffer (10mM Tris-HCl pH 7.5, 10mM NaCl, 3mM MgCl_2,_ 1%BSA, 0.1% NP-40, 0.1% Tween-20, 0.01% digitonin)^59^. If nuclei were processed fresh, OmniATAC lysis buffer was supplemented with Pitstop2 (70mM, Abcam 120687) to increase nucleus permeability to Tn5^60^. Nuclei were gently isolated and permeabilized by Dounce homogenization and mechanical pipetting for a maximum of 3min in cold conditions. 1.7mL of cold ATAC wash buffer (10mM Tris-HCl pH 7.5, 10mM NaCl, 3mM MgCl_2,_ 2%BSA, 0.1% Tween-20) was added and the nuclei filtered through 40mm strainer into a new 2mL LoBind tube. Nuclei were pelleted at 500g in a swinging bucked rotor, 7min at 4°C. The resulting pellet was washed twice in cold PBS-1%BSA, gently resuspended in 1x Diluted buffer (10x genomics) and filtered through 40mm cell strainer (Flowmi^R^).

If nuclei suspension was purified from debris and aggregates using FACS (see below), nuclei were Dounce homogenized in OmniATAC lysis buffer without digitonin and mildly fixed in 0.1% methanol-free formaldehyde (PFA) (ThermoFisher 28906) to mitigate nuclei damage during sorting. Briefly, after washing in ATAC wash buffer, nuclei were incubated in PBS-1%BSA 5min on ice, gently resuspended, and fixed 5min at RT by adding 1%PFA in PBS to reach a final concentration of 0.1%PFA. Reaction was quenched by adding glycine (0.125M final), Tris-HCl pH 8 (50mM final) and BSA (1.7% final), 5min at 4°C. Nuclei were pelleted at 500g, 5min at 4°C, and washed once with cold PBS-1%BSA. The resulting pellet was gently resuspended and stained in PBS-1%BSA with DAPI (1mg/mL final concentration) prior to FACS. One million of single nuclei (2n and 4n DNA content) were sorted using FACS Influx (100um nozzle, 12psi, cold conditions) into PBS-1%BSA. Nuclei were pelleted and permeabilized in 0.1x OmniATAC lysis buffer (10mM Tris-HCl pH 7.5, 10mM NaCl, 3mM MgCl_2_, 1%BSA, 0.01% NP-40, 0.01% Tween-20, 0.001% digitonin) supplemented with 70mM Pitstop2, 2min on ice while gently pipetting. After washing, nuclei were processed as described above for fresh nuclei (sample name: 2_Gastrula_fix).

Prior to each Chromium scATAC-seq run (10x Genomics), an aliquot of nuclei suspension was taken to assess their quality and concentration. For this, nuclei were stained with DAPI and loaded on a Neubauer chamber for counting under a fluorescence microscope. Nuclei concentration was adjusted to encapsulate ∼10k nuclei from each sample with the 10x Chromium platform after tagmentation in bulk. scATAC-seq libraries from gastrula stage were prepared using the Chromium scATAC v2 (Next GEM) kit from 10x Genomics, following the manufacturer’s instructions.

*<u>2) Whole adult scATAC-seq</u>*. 2/3 months old *Nematostella* polyps were obtained from non-sexed wild-type polyps, starved at least 3 days, and spawned one day before dissociation to avoid any possible contamination with gametes. 2-4 adult polyps were washed in PBS before socking them into ice-cold TST lysis buffer (10mM Tris-HCl pH 7.5, 146mM NaCl, 1mM CaCl_2_, 21mM MgCl_2_, 0.03% Tween-20, 1x cOmplete protease inhibitor)^61^. Polyps were transferred on a clean slide on ice and minced with a pre-chilled knife into small chunks. Chopped tissue was then gently crushed in an ice-cold Dounce homogenizer until homogenous suspension was achieved, and further dissociated by pipetting (p1000 strokes). Sample was maximum 12min in TST lysis buffer, then diluted with 1 volume of cold 2%BSA in ST buffer (without tween-20). Resulting cell/nuclei suspension was filtered through 70mm strainer into LoBind protein tube and pelleted at 800g, 5min at 4°C. To purify single nuclei from debris and aggregates, sample was mildly fixed in 0.1% PFA prior to FACSorting as described above. Between 700k-1M single nuclei were sorted into PBS-2%BSA, pelleted and permeabilized 2min in cold 0.1x OmniATAC lysis buffer with Pitstop2. When nuclei from adult samples were processed fresh (without PFA fixation), NP-40 was added to TST lysis buffer (0.01% NP-40 final concentration) after Dounce homogenization and further dissociated 5min by pipetting. In this case, nuclei were purified from debris using an OptiPrep® continuous density gradient. Fresh purified nuclei were permeabilized in ice-cold 1x OmniATAC lysis buffer with Pitstop2 for 4min while gently pipetting. Finally, fresh or fixed and permeabilized nuclei were washed in ATAC wash buffer, resuspended in 1x Diluted buffer and filtered through 40mm strainer (Flowmi) before counting.

16 scATAC-seq libraries were generated from adult fixed samples (sample name: 3-15 & 17-19 Adult_Fix), and 1 scATAC-seq library from fresh sample (16_Adult_Fresh). All of them using the Chromium scATAC v1.1 (Next GEM) kit from 10x Genomics and following the manufacturer’s instructions.

*<u>3) scATAC-seq of NvElav1::mOrange-positive cells</u>.* To enrich our adult scATAC-seq dataset with neural cells, *NvElav1::mOrange*-positive cells were purified by FACS as previously described^62^, with minor modifications. Briefly, one month old *NvElav1::mOrange* positive polyps were dissociated at 25°C in calcium- and magnesium-free NM (CMF/NM) containing 5mM EDTA and 0.25% α-chymotrypsin (Sigma, C4129). Single cell suspensions were then stained with Hoechst 33342 (1µg/ml, ThermoFisher 62249) and TO-PRO-3 (50nM, invitrogen T3605) to remove debris and non-viable cells by FACS (FACS Aria II, 100um nozzle, cold conditions). Nuclei from 150k sorted mOrange-positive cells were isolated, mildly fixed in 0.1%PFA and permeabilized (samples: 20&21 Elav_fix), or directly permeabilized in OmniATAC lysis buffer with Pitstop2 (samples: 22-24 Elav_fresh). Finally, permeabilized nuclei were washed in ATAC wash buffer, resuspended in 1x Diluted buffer and encapsulated using 10x Chromium platform. Five scATAC-seq libraries were generated using the Chromium scATAC v1.1 (Next GEM) kit from 10x Genomics, following the manufacturer’s instructions.

*<u>4) Whole-adult scMultiome (ATAC+RNA).</u>* Two adult wild-type polyps were dissociated and stained for FACSorting as described above for *NvElav1::mOrange* samples. Nuclei from 250k single viable cells were isolated and permeabilized for 3 min in ice-cold 0.1x OmniATAC lysis buffer supplemented with Pitstop (70mM), RNAse inhibitor (1U/ul) and DTT (1mM). Nuclei were then washed in ATAC wash buffer and resuspended in 1x Diluted Nuclei Buffer, both supplemented with RNAse inhibitor (1U/ul) and DTT (1mM). Nuclei were counted after filtering through 40mm strainer with the help of DAPI, and encapsulated using 10x Chromium platform.

One run of Chromium Next GEM single cell multiome kit from 10x Genomics was performed, following the manufacturer’s instructions and performing 8 PCR cycles in Step 5.1 for scATAC library construction (24_Adult_Fresh_MultiomeATAC), or 9 PCR cycles of cDNA amplification in Step 6.1 for scRNA library construction (24_Adult_Fresh_MultiomeGE).

*<u>5) Whole-adult 5’ scRNA-seq</u>*. Single cell suspensions were obtained after ACMEsorbitol (0.4M) fixation and dissociation as previously described^63,64^. One single-cell 5’ GE library was generated using the Chromium Next GEM 5’ GEX scRNAseq v2 kit from 10x Genomics, following the manufacturer’s instructions with 14 PCR cycles of cDNA amplification.

scATAC-seq libraries (**Supplementary Table 1**) were sequenced to reach ∼5,000 reads per cell (average: 19,575), with median 2,788 unique fragments per cell on average. (50/8/16/50). The scATAC-seq library derived from scMultiome kit (07564AAD) was sequenced at 3,929 reads per cell and 1,556 unique fragments per cell (50/8/24/49), while the scRNAseq library (07563AAD) was sequenced at 8,080 reads per cell and median 789 UMIs per cell (28/10/10/90). The 5’ scRNAseq library (07575AAD) was sequenced at 24,297 reads per cell and median 863 UMIs per cell (26/10/10/90). All libraries were sequenced using Illumina NextSeq500 platform.

### scATAC-seq processing

We processed scATAC sequencing data using a modified scATAC-pro workflow^65^. Briefly, we mapped sequencing reads to *Nematostella* Darwin Tree of Life genome^66^ using bwa^67^, and filtered nucleosome free reads for downstream analysis. Initial cell calling with was done using EmptyDrops^68^, with false discovery rate 0.05. scATAC downstream analysis was done using ArchR^69^. Cells called with EmptyDrops that had TSS enrichment below 4 and less than 200 fragments were filtered out. We also added doublet scores using ArchR’s in silico doublets method, and removed cells predicted to be doublets using filterRatio = 1 (4% of input cells). We then performed dimensionality reduction using iterative LSI (4 iterations) and clustering using top 10,000 variable features, with resolution 0.3. We identified and removed clusters of low-quality cells with TSSEnrichment < 8. We then repeated dimensionality reduction and clustering iteratively until all resulting clusters were of good quality. Next we used SEACells^70^ for grouping cells into metacells, with target of ∼75 single cells per metacell. Metacells obtained from SEACells approach were grouped in clusters and broadly annotated by label transfer from scRNA-seq data using AUCell^71^. Additional, more specific cell type annotations were then assigned by inspecting the accessibility (gene scores, described below) of known marker genes. Final consensus set of peaks was generated by peak calling for pseudobulk aggregated cell types using MACS2^72^ and iterative reduction approach implemented in ArchR.

### Peaks to gene assignment and gene score calculation

To each gene we assigned peaks that are within the gene’s body or <10 kb away from the gene’s TSS, unless they were coming after (upstream or downstream) a TSS of another gene (implemented in *mta_match_peaks_to_genes()* function). Initially, 52,526 (63%) peaks were assigned to a single gene, and 31,098 (37%) peaks were assigned to more than one gene. For the latter ones, we refined the assignment by taking into account peaks co-accessibility (calculated by Cicero) and the correlation of accessibility to gene expression. Briefly, for all co-accessible peaks groups (co-accessibility > 0.5) assigned to more than one gene, we looked for a sharp drop in ranked peak-to-gene correlation (Δcorrelation < -0.1) for all peaks in the group, and removed those assignments that followed the drop (this procedure is implemented in *mta_refine_peaks_to_genes_by_coaccessibility()* function). As a result, we refined the assignment of 2,142 peaks. Next, we calculated gene scores as a weighted sum of the accessibility of all peaks assigned to gene. Each peak is weighted by distance from the gene (peaks inside the gene body get maximum weight of 1) and by peak specificity, measured by Gini index (**Figure 1D**). This procedure is implemented in *mta_gene_scores()* function. Using 5’ scRNA-seq and H3K4me3 data together with scATAC peaks, we devised a decision tree approach (**Figure S3A**) to assign promoters to genes, and further classify them as constitutive promoters (CP) which are accessible in all cell types, specific promoters (SP) accessible in one or multiple cell types, but not all, and potential alternative promoters (AP), with different promoters being used in different cell types (this is implemented in *mta_class_promoters()* function).

### Motif archetypes

We aimed to collect a comprehensive catalogue of all possible TF binding motifs in *Nematostella* genome. To this end, we combined motifs for *Nematostella* TFs that were either determined experimentally or inferred from other species based on TFs’ DNA-binding domain (DBD) sequence similarity, with motifs we found to be significantly enriched or depleted in either all accessible or specifically accessible peaks in cell types, or enriched in different promoter classes (AP, SP, CP). To reduce redundancy of this comprehensive catalogue of motifs, we calculated pairwise similarities between PWMs using *compare_motifs()* function from universalmotif R package (Pearson correlation coefficient (PCC) with normalise.score option to favour alignments which leave fewer unaligned positions, as well as alignments between motifs of similar length), and then we applied complete hierarchical clustering, choosing the number of clusters that maximises the ratio of within- and between-cluster median pairwise similarities. These initial clusters of similar motifs were further split into smaller clusters that contain only motifs above desired similarity threshold (0.8). For all the motifs in each cluster we applied IC block filtering^73^, retaining only motifs with a block of at least 4 consecutive bases with IC >= 0.5 (ungapped motif), or at least two blocks of at least 3 consecutive bases with IC >= 0.5 (gapped motif). Then we generated a consensus PWM by averaging aligned PWMs at each position. Finally, we trimmed off the leading and trailing positions with IC < 0.5 in the consensus archetype motif. This entire procedure is implemented in the mta_merge_archetype() function. By doing this, we reduced the filtered input set of 2,951 motifs to 1,292 archetypes (**Extended Data** Fig. 6). We show that min-max normalised motif scores in accessible peaks are comparable for archetypes and highest scoring motifs in each archetyping cluster, as well as the motif enrichments in cell type specific peaks (**Extended Data** Fig. 6g).

### Assigning motifs to TFs

Binding motifs have been determined experimentally only for a subset of *Nematostella* TFs^45^. We devised a computational approach to assign a motif from our comprehensive set of motif archetypes (from now on, motifs) to each TF gene without experimentally determined motif (**Extended Data** Fig. 4b). We first calculated motif activity scores for all motifs using chromVAR^74^. Next, we calculated correlations between each motif’s activity score and both expression and accessibility (i.e. gene score) of each TF. We ranked motifs based on gene score correlation and for each gene we selected the best correlated motif of the same structural class, if either expression or gene score correlation was greater than 0.3. To improve the accuracy of assignment, particularly for large structural classes such as Homeodomains, we also considered closest human, mouse, rat and zebrafish orthologs of each TF, and if the motif activity of ortholog gene’s motif correlated better than that of previously selected archetype, we assigned that ortholog motif to a given *Nematostella* TF (**Extended Data** Fig. 4c, d).

### Gene Regulatory Network inference

We used *in silico* ChIP method^75^ to link TFs to target scATAC peaks. Briefly, *in silico* ChIP links TFs to a peak, if the peak contains a motif hit for the TF and if the accessibility of the peak correlates with the RNA expression of the TF. Correlation between peak accessibility and RNA expression at metacell level was calculated after mapping each scATAC metacell to the best-correlated scRNA metacell of the same broad cell type. Motif hits were determined using *findMotifHits()* function from monaLisa R package ^76^ with 95th quantile of genome-wide motif scores distribution for each motif used as a minimum score for counting a hit. *in silico* ChIP outputs a matrix of TF binding scores for each peak, ranging from 0 to 1, and it is necessary to select a threshold value for +. For each motif we calculated its cell type activity as a Z-score of accessibility deviation of the target peaks set (selected with different *in silico* ChIP binding score thresholds) in a given cell type, compared to assumption of equal chromatin accessibility across cell types, and normalised by a set of background peaks matched for GC and average accessibility. From this, we selected 0.1 as a binding score cut-off because this was the value that maximised the correlation of TF expression and TF activity for the majority of TFs.

TFs and target peaks for which binding score is > 0.1 constitute a global GRN. We further partitioned this into cell type specific GRNs by filtering TFs based on expression and TF activity, and filtering target genes based on expression and accessibility. We used a per-cell type 0.4 quantile threshold of expression fold change to filter genes (both TFs and target genes) by expression. To filter peaks, we used a per-cell type 0.4 quantile threshold of normalised peak accessibility. To filter TFs based on activity, we used a Z score threshold of 4. For plotting GRNs (**Fig. 3i**, **Extended Data** Fig. 8), we additionally filtered out genes with expression fold change < 1.2.

### Sequence models

We trained per-cell type gapped kmer classifiers with gkm-SVM^77^ on the set of accessible peaks in each cell type (Log2FC > 1 and p-value < 0.1), with 70-30 train-test split and 5-fold cross validation. Of note, we used a relaxed threshold for selecting specific peaks, because we wanted the models to learn more general CRE sequence features that might be shared across similar cell types. We applied each model to all left-out sets of peaks and calculated test-set AUC statistics. We used gkmexplain^41^ on top 1,000 scored peaks per model to identify important sequence features for each cell type classifier. We also trained two types of deep learning models on *Nematostella* chromatin accessibility data. We trained ChromBPNet v1.5 models^78^ that predict accessibility at base pair resolution from underlying CRE sequences. We first trained bias models to learn Tn5 sequence biases to be regressed out during the actual models training. Then we trained models to predict total counts and accessibility profile signals for each cell type using cell type accessible peak and GC-matched non-peak sequences as inputs. The models were trained using default architcture with 4 dilation layers and 512 filters, on 500 bp sequences as input, with prediction on 250 bp. We also trained the peak regression CREsted model (https://github.com/aertslab/CREsted), which adapts the original ChromBPNet architecture. CREsted model was trained on peak logcounts, normalized by subtracting mean value across class. The model was trained using the default architecture, with learning rate 5e-1 and Huber loss function, on 500 bp sequences as input, with prediction also on 500 bp. During training of both CREsted and chromBPNet models, we left out CREs on one chromosome for validation (NC_064034.1) and on another for testing (NC_064035.1). We used SHAP DeepExplainer^79^ to estimate the predictive importance of each base in CRE sequence, and TF MoDISco-lite^80^ to identify sequence patterns (i.e. motifs) that are relevant for accessibility prediction. As input for TF MoDISco, we selected 5000 most specific regions per class, and out of those used the 1000 regions with the highest predictions scores for that class. We then used the same archetyping procedure as described above to reduce redundant patterns from different models, with the only difference being that here we used Jannson-Shannon divergence (JSD) as a metric of motif similarity. To compare pattern archetypes to known motif archetypes, we calculated JSD for every motif archetype-pattern archetype pair in two ways: along the entire length of motifs alignments (JSD_complete_) and only along the overlapping fraction of alignment (JSD_min_) - in this way we could better distinguish novel motifs from similar motifs in different contexts (**Extended Data** Fig. 6h).

### Generation of *Nematostella* transgenic lines

*NvGabra2::mOrange* transgenic reporter lines driven by differentially accessible alternative promoters in tentacle retractor muscle (tRM-AP) or neuron Pou4/FoxL2 (NeuroPou4/FoxL2-AP) cells were generated by meganuclease-mediated transgenesis as described by Renfer and Technau^81^.

The genomic coordinates for the ca. 2.8kb regulatory region of tRM-AP are 11660621-11657766 on minus strand chromosome 2. The genomic coordinates for the ca. 2kb regulatory region of NeuroPou4/FoxL2-AP are 11644315-11642257 on the same minus strand of chromosome 2^66^. These regulatory regions were cloned in frame with mOrange reporter gene into the meganuclease (I-Sce1)-mediated transgenesis vector kindly provided by Technau lab ^81^. Wild-type fertilized eggs were injected with a mix containing: plasmid DNA (20 ng/µl), I-Sce1 (1 U/µl, NEB R0694), Dextran Alexa Fluor^TM^ 488 (50 ng/µl, Life Technologies D22910) and CutSmart buffer (1x). The mix was incubated at 37 °C for at least 20 min, then injection was performed at 18°C with a FemtoJet® 4i microinjector (Eppendorf). Constructs and/or transgenic lines are available from the authors upon request.

### Immunofluorescence

One month old F1 polyps derived from *NeuroPou4/FoxL2-AP::mOrange* transgenic line were relaxed in 0,34% MgCl_2_/NM solution to prevent tentacle contraction before cutting with a sharp knife at the level of the pharynx. The resulting heads were fixed in 3,7% formaldehyde in PBS-0.1%Tween20 (PBTw) O/N at 4 °C, washed several times in PBTw the day after, and left in PBS O/N at 4 °C.

For immunostaining against mOrange, samples were washed several times in PBS-0.3%TritonX (PBTx) during 1h at room temperature (RT), blocked in blocking solution (1% BSA/5% Normal Goat Serum/PBTx) for 1h at RT, and incubated with rabbit anti-DsRed primary antibody (1:100, Clontech 632496) in blocking solution O/N at 4 °C. Samples were then washed several times in PBTx-BSA during 2h at RT, blocked in blocking solution for 30min at RT and incubated with goat anti-rabbit Alexa568 secondary antibody (1:250, Life Technologies A11011) in blocking solution O/N at 4 °C. Samples were then washed 5x in PBTx-BSA during 2h at RT, 5min in PBS, and left in 70% Glycerol in PBS at 4 °C for at least O/N. Samples were mounted in ProLong^TM^ Glass antifade mountant (Thermo Fisher Scientific, P36982) and imaged on a Leica SP8 confocal microscope. Images were extracted from z-stacks with Fiji and adjusted for brightness/contrast applied to the whole image.

### Live Imaging

Adult F0 polyps derived from *tRM-AP::mOrange* transgenic cassette insertion were relaxed in 0,34% MgCl_2_/NM solution before cutting with a sharp knife at the level of the pharynx. The resulting heads were then mounted in a slide with 2,43% MgCl_2_/NM solution for live imaging on a Leica SP8 confocal microscope and images extracted as described above.

---

**Extended Data Figure 1.**
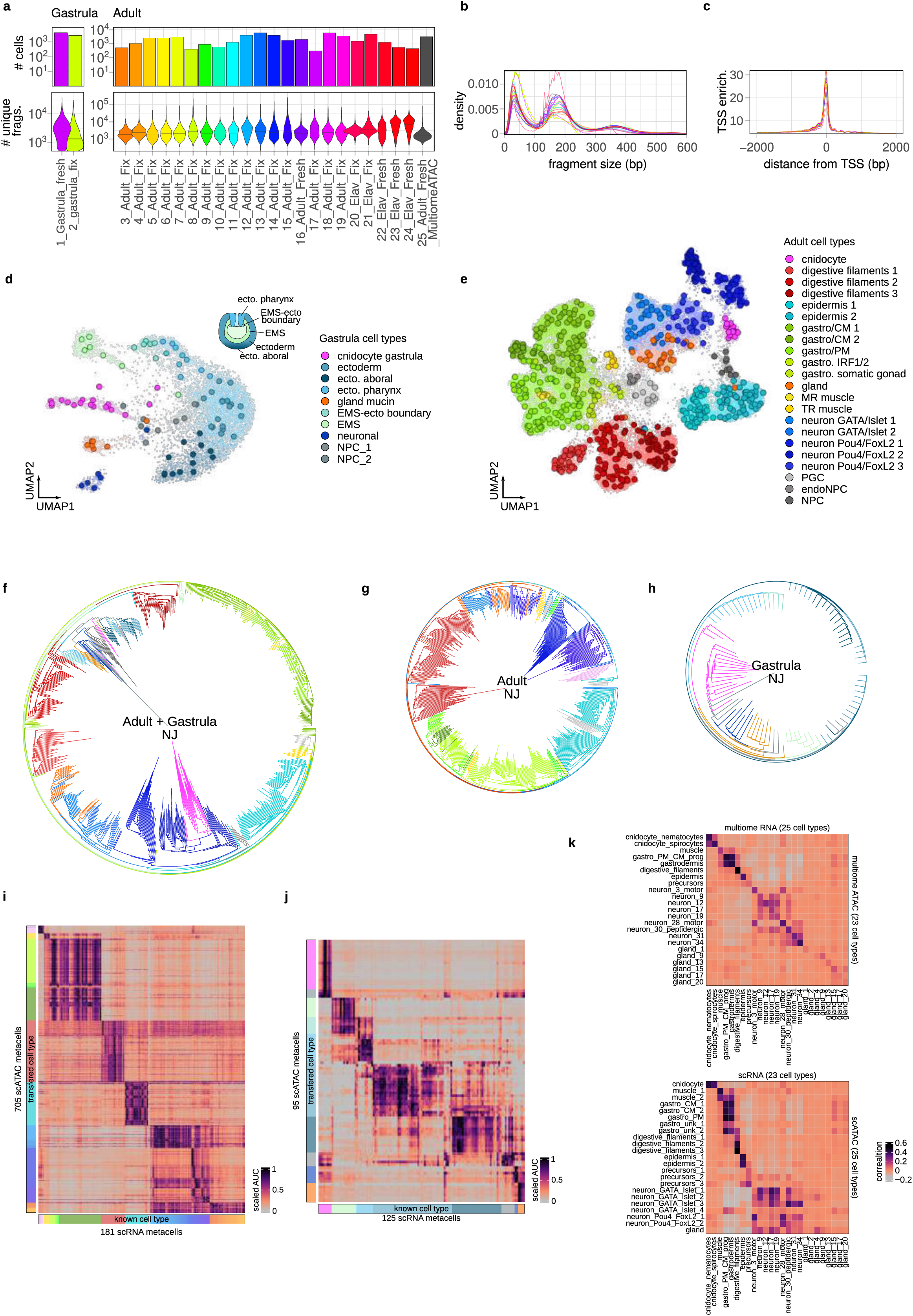
scATAC-seq dataset QC, clustering and annotation. **a,** Number of cells (top) and unique fragments per cell (bottom), **b,** scATAC-seq fragment size distribution for each sample. **c,** TSS enrichment signal for each sample. **d,** UMAP projection of single cells and metacells for gastrula dataset. **e**, UMAP projection of single cells and metacells for adult dataset. **f**, NJ clustering of metacells for adult and gastrula together, only for adult (**g**) and only for gastrula (**h**). **i**, Annotation transfer heatmap for adult scATAC-seq clusters. **j**, Annotation transfer heatmap for gastrula scATAC-seq clusters. **k**, Comparison of ATAC and RNA correlations for multiome (top) and separately profiled scATAC-seq and scRNA-seq data (bottom).

**Extended Data Figure 2.**
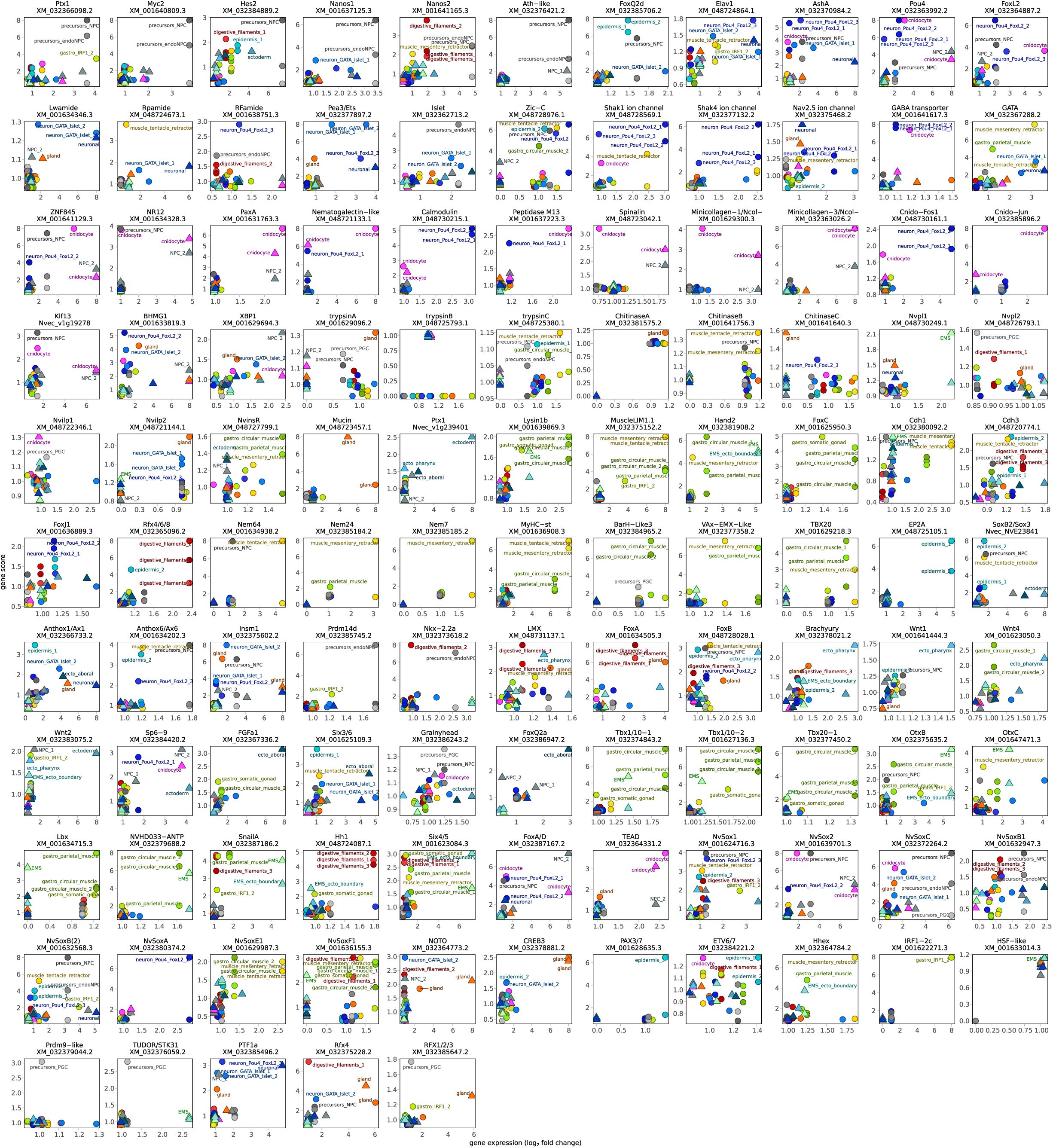
Comparison between accessibility scores and expression for selected marker genes.

**Extended Data Figure 3.**
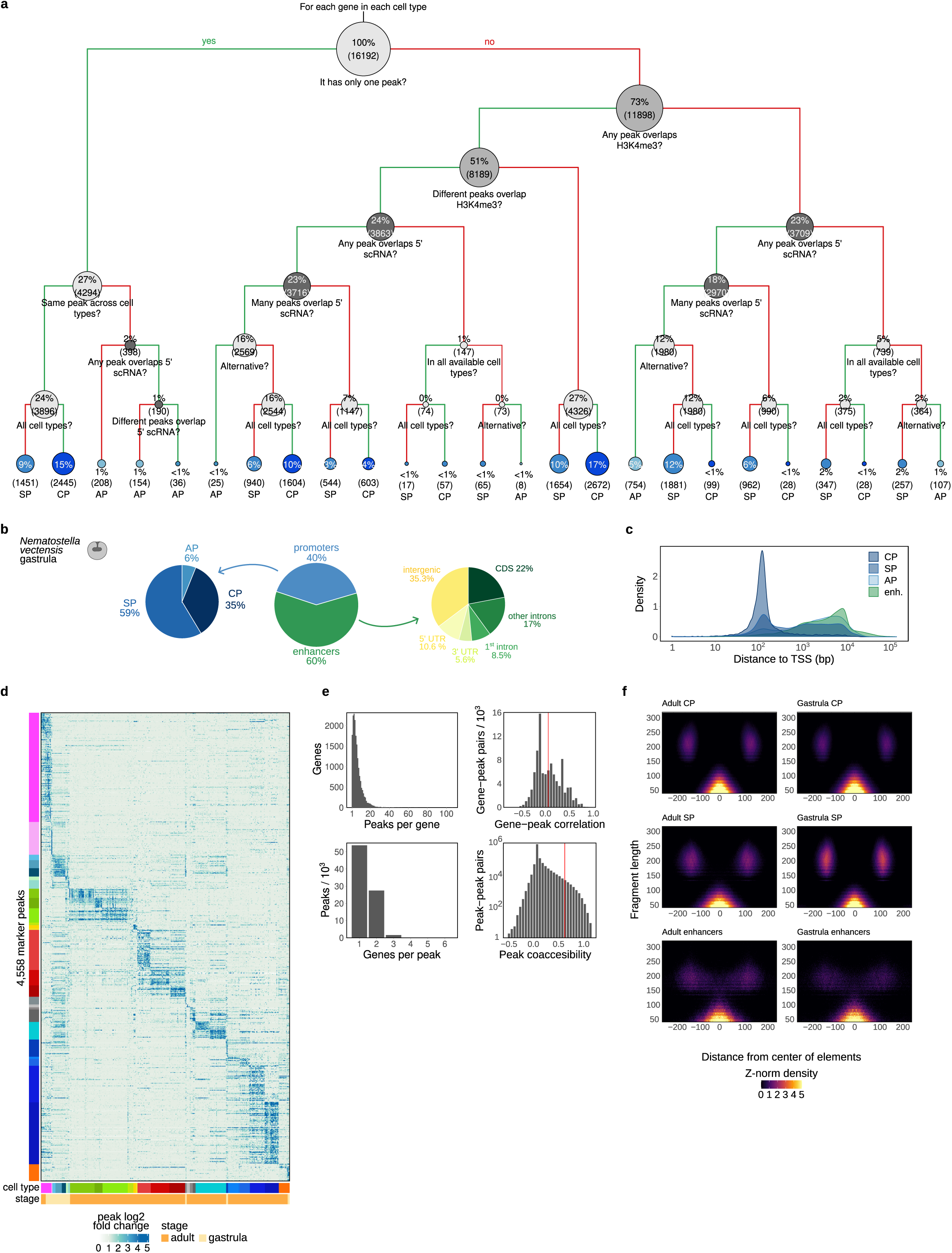
*Cis*-regulatory element classification. **a**, Decision tree used to classify CRE into different promoter types. **b**, Fraction of gastrula CREs classified as promoters and enhancers. Promoters are further classified as constitutive promoters (CP), specific promoters (SP) and alternative promoters (AP). Enhancers are classified based on their overlap with different genomic regions. **c**, Distance to the nearest TSS distributions for different types of CREs. **d**, Heatmap of peak accessibility per cell type, cell types color-coded as in Fig.1. **e**, Summary peak statistics. Number of peaks per gene (top-left) and number of genes that each peak gets assigned to (bottom-left), correlation across metacells between peak accessibility and expression of the genes they are assigned to (top-right), and co-accessibility across cell clusters of all pairs of peaks (bottom-right). **f**, V-plots showing tagmentation fragment size distributions (y-axis) at different distances (x-axis) around CP, SP and distal CREs in adult and gastrula stages.

**Extended Data Figure 4.**
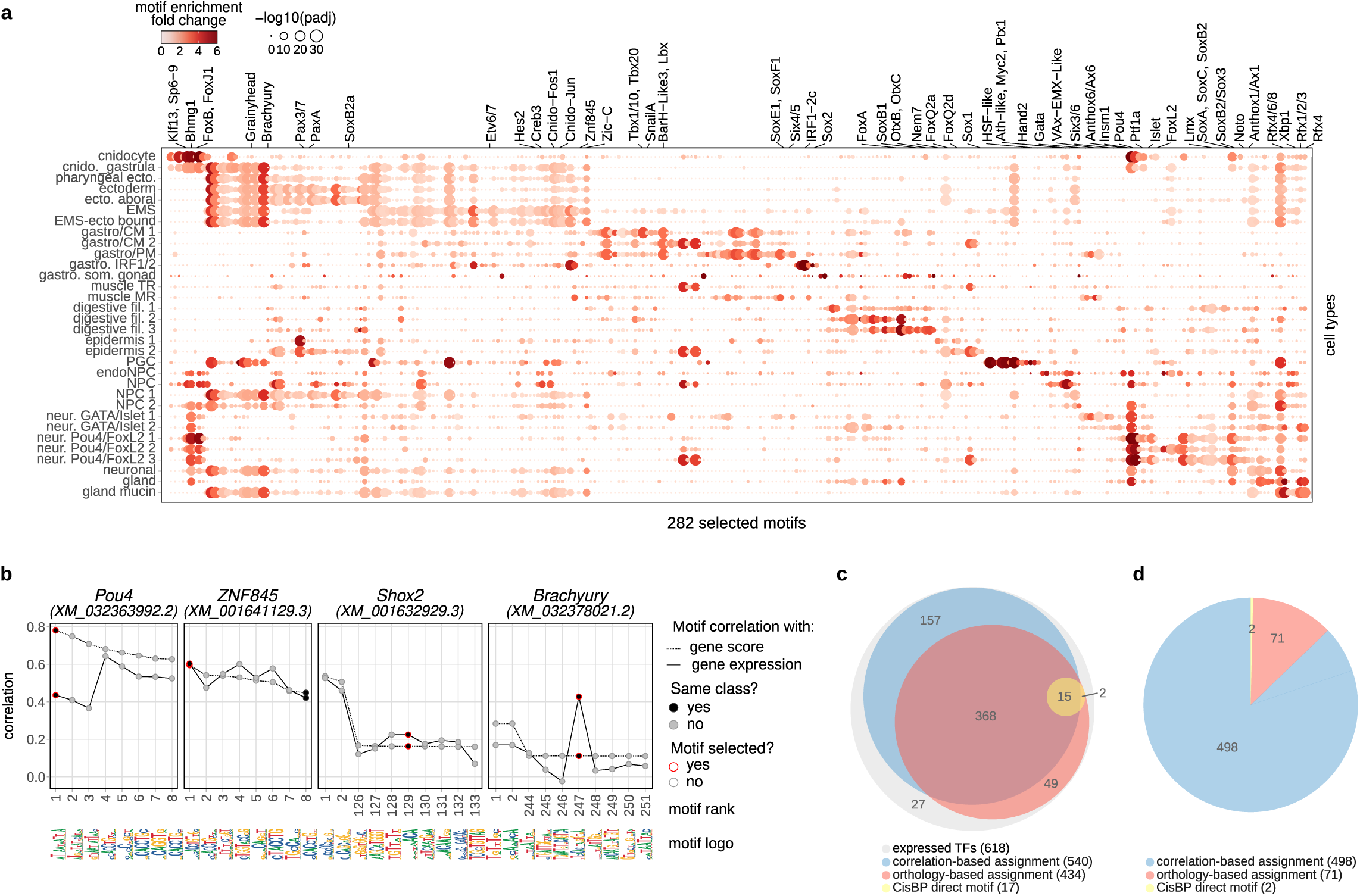
Motif enrichment analysis and assignment to TFs. **a**, Dotmap of representative motifs enriched in different cell types. Significantly enriched motifs (fold change > 1, padj < 0.001) are grouped by motif-motif similarity and top enriched representative motif for each cluster are indicated. **b**, Correlation-based approach for assigning motifs to TFs. For each TF, we rank motifs based on correlation of motif activity to TF accessibility and expresssion, and assign it top ranking motif of the same structural class. **c**, Euler diagram showing TF coverage using different motif-to-TF assignment methods for expressed *Nematostella* TFs. **d**, Final motif assignment sources for expressed *Nematostella* TFs.

**Extended Data Figure 5.**
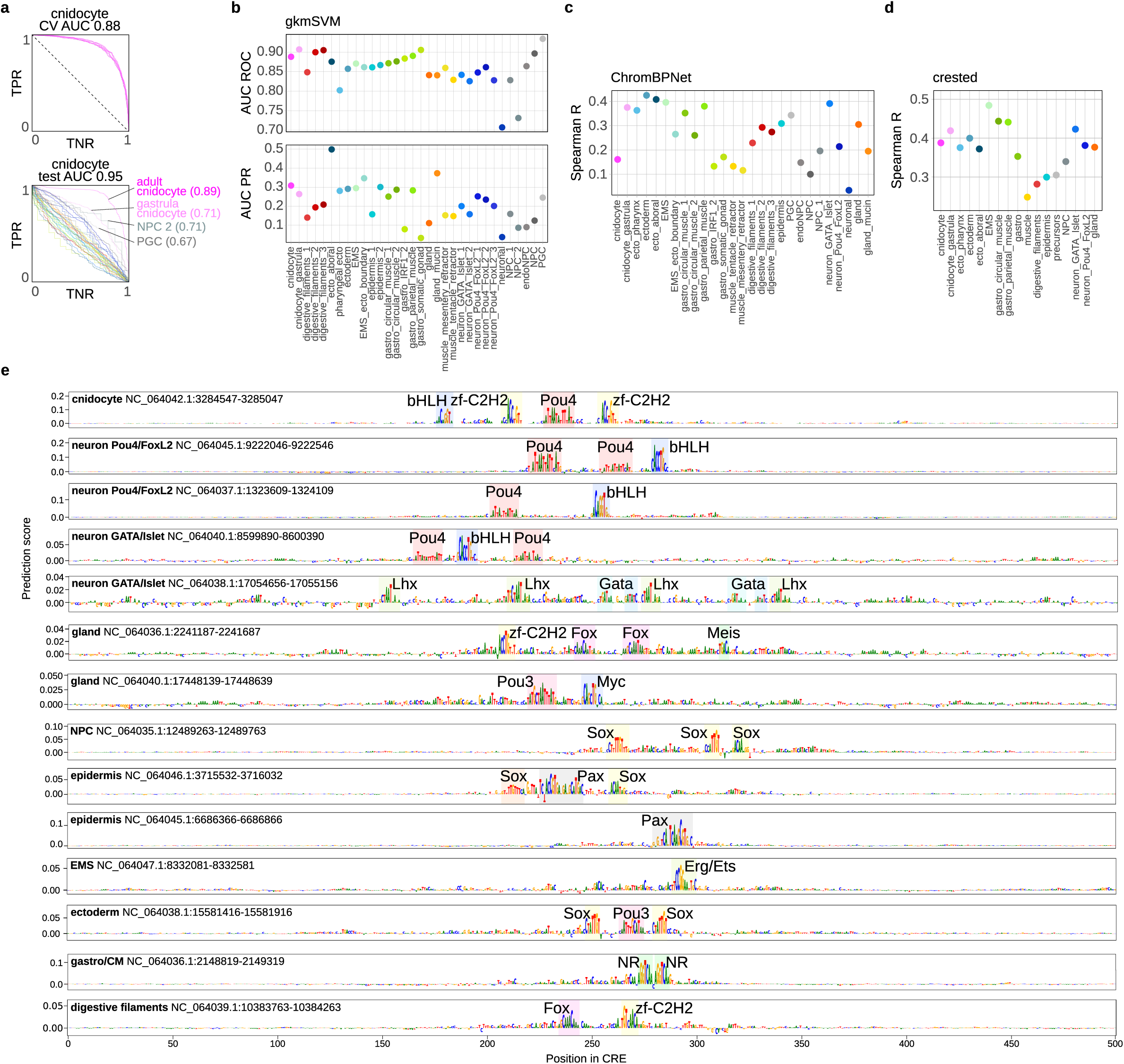
Sequence models. **a**, Five-fold cross-validation (CV) area under the curve (AUC, top) and test set AUC (bottom) for gkm-SVM classifiers trained on adult cnidocytes. **b**, Area under receiver operator curve (AUC ROC, top) and area under precision recall curve (AUC PR, bottom) for all cell type gkm-SVM classifiers. **c**, Spearman correlation for ChromBPNet predicted accessibility counts in test set peaks. **d**, Spearman correlation for crested predicted accessibility counts in test set peaks. **e**, Nucleotide importance scores for top scored CREs in different cell types.

**Extended Data Figure 6.**
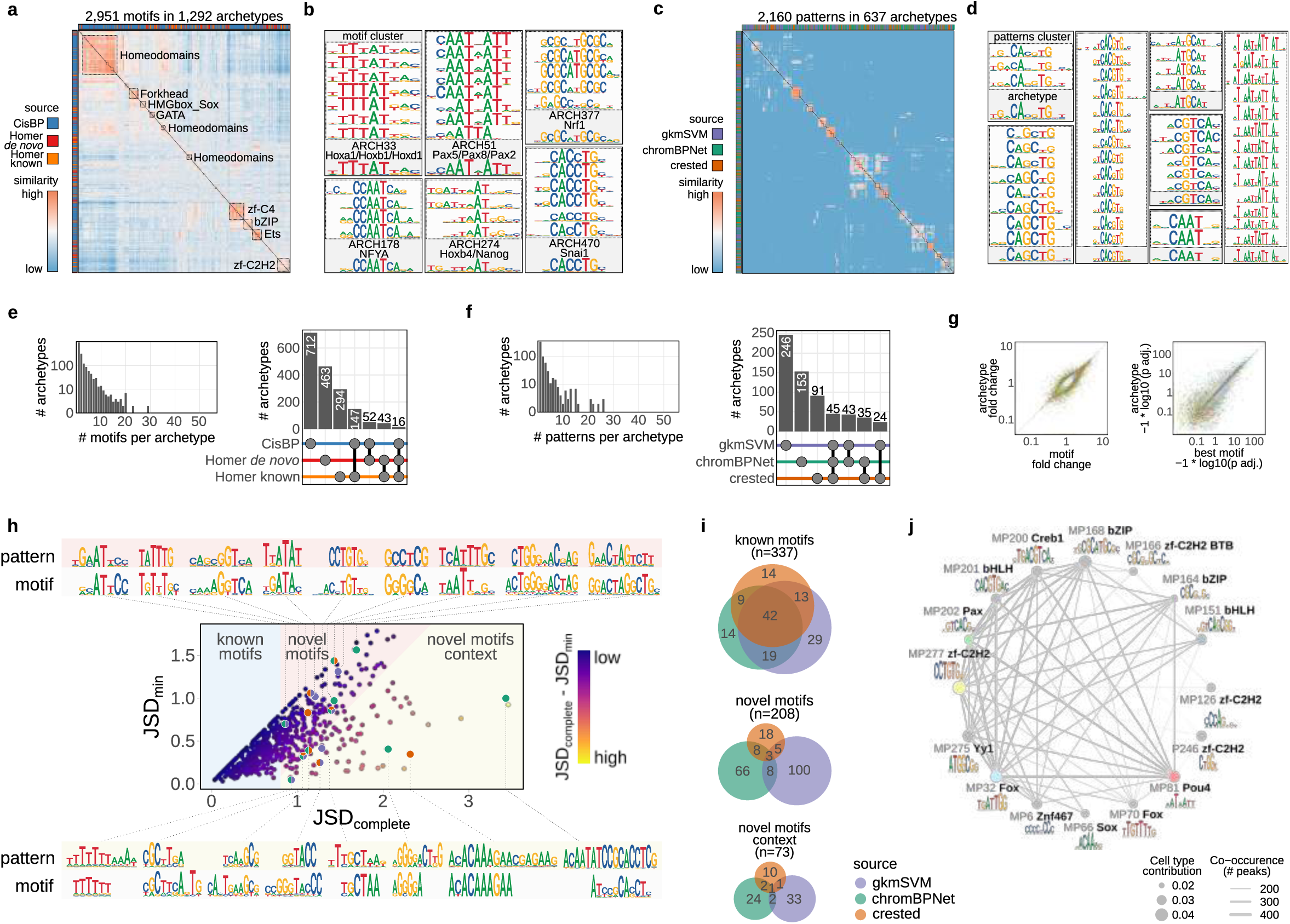
Sequence motif discovery. **a,** Heatmap showing pairwise motif similarities used to generate motif archetypes from enriched motifs. **b,** Examples of motif clusters. **c,** Examples of motif archetypes. **d**, Same as (a) and (b) for patterns discovered with sequence models. **e**, Number of motifs per archetype (left) and number of archetypes composed of motifs from different sources (right). **f**, Number of patterns per archetype (left) and number of archetypes composed of patterns from different sequence models (right). **g**, Comparison of motif enrichment fold change (left) and adjusted p-value (right) for archetypes versus best scoring motif in each archetype cluster. **h**, For all pattern archetypes and their most similar motif archetype, Jensen-Shannon divergence (JSD) calculated across the best pairwise alignment of archetypes (x-axis, JSD_complete_), and calculated across the best alignment spanning the length of shorter archetype (y-axis, JSD_min_). Based on these two metrics, pattern archetypes are classified as being novel motifs, having novel context or resembling known motifs from motif enrichment analysis. **i**, Euler diagrams summarizing the source of pattern archetypes for each of these three categories. **j,** Co-occurence network of pattern archetypes with contribution in cnidocytes. Size of the node reflects its cell type contribution, and width of the connection scales with the number of CREs in which two motifs co-occur.

**Extended Data Figure 7.**
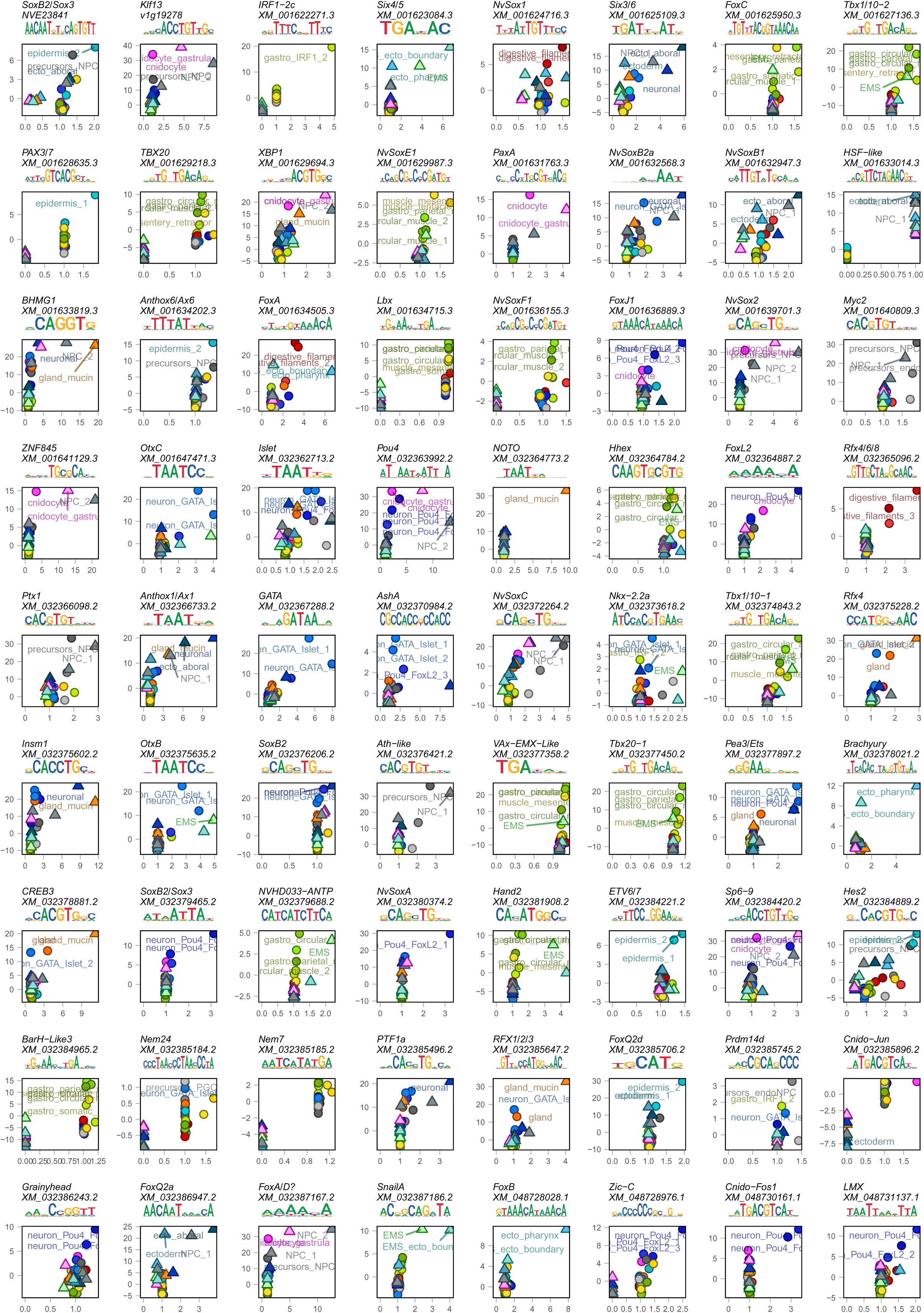
Examples of TF expression and TF motif activity correlations for selected marker genes.

**Extended Data Figure 8.**
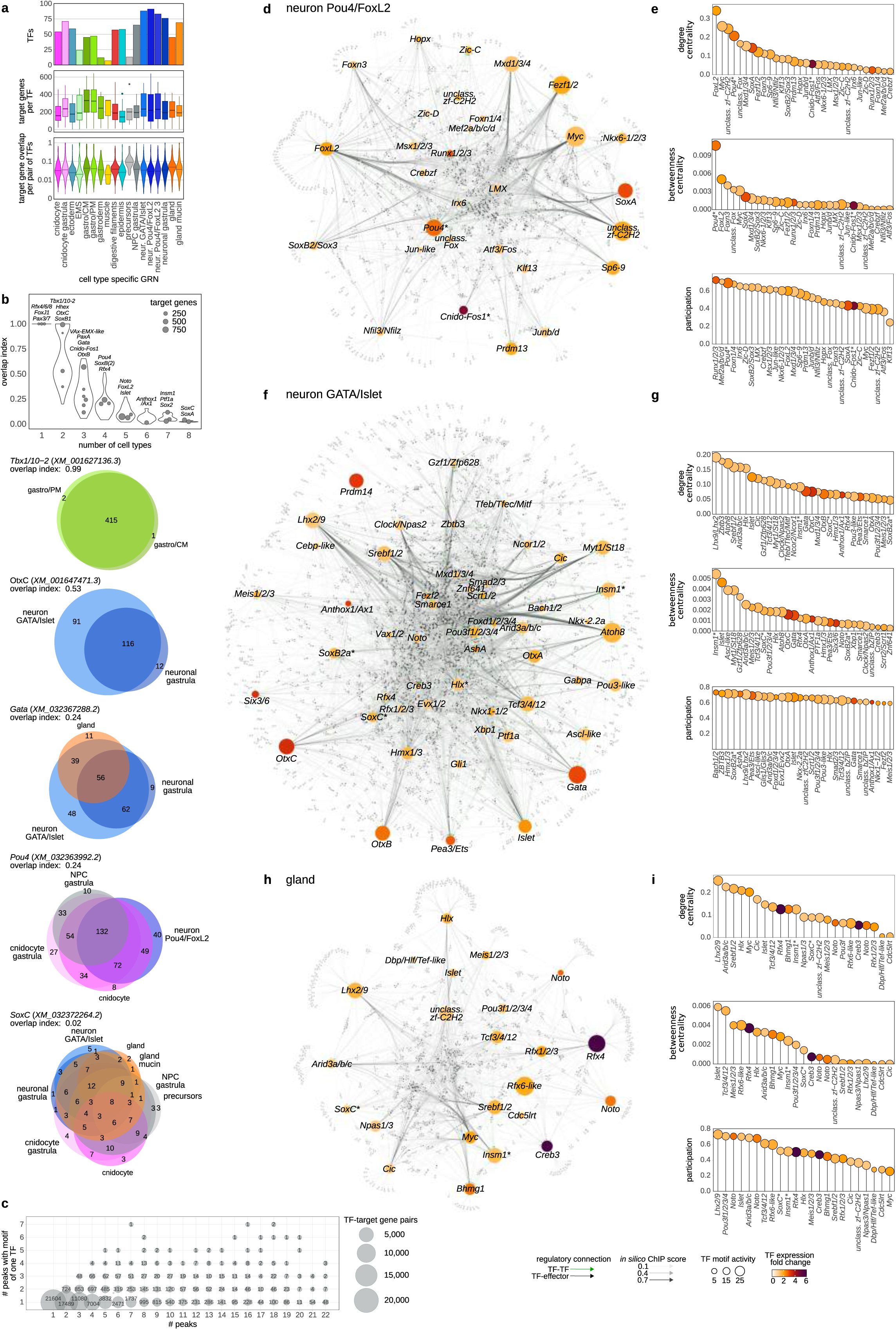
Cell type gene regulatory networks. **a**, Number of TFs in GRNs inferred for each broad cell type (top), number of genes targeted by each TF (middle), and fraction of overlapping target genes for each pair of TFs (bottom). **b**, Overlap of target genes for the same TF across cell types, plotted for groups of TFs active in different number of cell types. Selected TFs are highlighted on the plot and overlap of their target genes is shown as Euler diagrams below. **c**, Number of CREs per target gene (x-axis) compared to number of CREs of the same gene with any single TF motif (y-axis). Most TFs have binding motif in a single CREs of their target genes**. d-g,** Additional inferred GRN and TF connectivity measurements for neuro-secretory cell types: GATA/Islet neurons (**d-e**), Pou4/FoxL2 neurons (**f-g**) and gland cells (**h-i**).

## References

1. Musser, J. M. et al. Profiling cellular diversity in sponges informs animal cell type and nervous system evolution. Science *(1979)* 374, 717–723 (2021).

2. Sebé-Pedrós, A. et al. Cnidarian Cell Type Diversity and Regulation Revealed by Whole-Organism Single-Cell RNA-Seq. Cell 173, 1520–1534.e20 (2018).

3. Sebé-Pedrós, A. et al. Early metazoan cell type diversity and the evolution of multicellular gene regulation. Nat Ecol Evol 2, 1176–1188 (2018).

4. Fincher, C. T., Wurtzel, O., de Hoog, T., Kravarik, K. M. & Reddien, P. W. Cell type transcriptome atlas for the planarian Schmidtea mediterranea. Science (1979) 360, eaaq1736 (2018).

5. Plass, M. et al. Cell type atlas and lineage tree of a whole complex animal by single-cell transcriptomics. Science (1979) 1723, eaaq1723 (2018).

6. Cao, J. et al. Comprehensive single-cell transcriptional profiling of a multicellular organism. Science (1979) 357, 661–667 (2017).

7. Levy, S. et al. A stony coral cell atlas illuminates the molecular and cellular basis of coral symbiosis, calcification, and immunity. Cell 184, 2973–2987.e18 (2021).

8. Najle, S. R. et al. Stepwise emergence of the neuronal gene expression program in early animal evolution. Cell 186, 4676–4693.e29 (2023).

9. Tanay, A. & Sebé-Pedrós, A. Evolutionary cell type mapping with single-cell genomics. Trends in Genetics 37, 919–932 (2021).

10. Janssens, J. et al. Decoding gene regulation in the fly brain. Nature 601, 630–636 (2022).

11. Cusanovich, D. a et al. Multiplex single-cell profiling of chromatin accessibility by combinatorial cellular indexing. Science *(1979)* 348, 910–914 (2015).

12. Calderon, D. et al. The continuum of Drosophila embryonic development at single-cell resolution. Science (1979) 377, (2022).

13. Domcke, S. et al. A human cell atlas of fetal chromatin accessibility. Science (1979) 370, eaba7612 (2020).

14. Sarropoulos, I. et al. Developmental and evolutionary dynamics of cis-regulatory elements in mouse cerebellar cells. Science (1979) 373, (2021).

15. Li, Y. E. et al. A comparative atlas of single-cell chromatin accessibility in the human brain. Science (1979) 382, eadf7044 (2023).

16. Zhang, K. et al. A single-cell atlas of chromatin accessibility in the human genome. Cell 184, 5985–6001.e19 (2021).

17. Cusanovich, D. A. et al. A Single-Cell Atlas of In Vivo Mammalian Chromatin Accessibility. Cell 174, 1309–1324.e18 (2018).

18. Minnoye, L. et al. Chromatin accessibility profiling methods. Nature Reviews Methods Primers 1, 10 (2021).

19. Schwaiger, M. et al. Evolutionary conservation of the eumetazoan gene regulatory landscape. Genome Res 24, 639–650 (2014).

20. Chari, T. et al. Whole-animal multiplexed single-cell RNA-seq reveals transcriptional shifts across Clytia medusa cell types. Sci Adv 7, 1–17 (2021).

21. Siebert, S. et al. Stem cell differentiation trajectories in Hydra resolved at single-cell resolution. Science (1979) 365, eaav9314 (2019).

22. Li, Y. et al. Single-cell transcriptomic analyses reveal the cellular and genetic basis of aquatic locomotion in scyphozoan jellyfish. bioRxiv 2023.02.06.527379 (2023) doi:10.1101/2023.02.06.527379.

23. Hu, M., Zheng, X., Fan, C.-M. & Zheng, Y. Lineage dynamics of the endosymbiotic cell type in the soft coral Xenia. Nature 582, 534–538 (2020).

24. Persad, S. et al. SEACells infers transcriptional and epigenomic cellular states from single-cell genomics data. Nat Biotechnol 2022.04.02.486748 (2023) doi:10.1038/s41587-023-01716-9.

25. Granja, J. M. et al. ArchR is a scalable software package for integrative single-cell chromatin accessibility analysis. Nat Genet 53, 935–935 (2021).

26. Steger, J. et al. Single-cell transcriptomics identifies conserved regulators of neuroglandular lineages. Cell Rep 40, 111370 (2022).

27. Richards, G. S. & Rentzsch, F. Regulation of Nematostella neural progenitors by SoxB, Notch and bHLH genes. Development 142, 3332–3342 (2015).

28. Lemaître, Q. I. B. et al. NvPrdm14d-expressing neural progenitor cells contribute to non-ectodermal neurogenesis in Nematostella vectensis. Nat Commun 14, 4854 (2023).

29. Cole, A. G. et al. Updated single cell reference atlas for the starlet anemone Nematostella vectensis. Front Zool 21, (2024).

30. Steinmetz, P. R. H., Aman, A., Kraus, J. E. M. & Technau, U. Gut-like ectodermal tissue in a sea anemone challenges germ layer homology. Nat Ecol Evol 1, 1535–1542 (2017).

31. Rentzsch, F., Fritzenwanker, J. H., Scholz, C. B. & Technau, U. FGF signalling controls formation of the apical sensory organ in the cnidarian Nematostella vectensis. Development 135, 1761–1769 (2008).

32. Lebedeva, T. et al. Cnidarian-bilaterian comparison reveals the ancestral regulatory logic of the β-catenin dependent axial patterning. Nat Commun 12, 4032 (2021).

33. Reddington, J. P. et al. Lineage-Resolved Enhancer and Promoter Usage during a Time Course of Embryogenesis. Dev Cell 55, 648–664.e9 (2020).

34. Weintraub, A. S. et al. YY1 Is a Structural Regulator of Enhancer-Promoter Loops. Cell 171, 1573–1588.e28 (2017).

35. Lenhard, B., Sandelin, A. & Carninci, P. Metazoan promoters: emerging characteristics and insights into transcriptional regulation. Nat Rev Genet 13, 233–45 (2012).

36. Haberle, V. & Lenhard, B. Promoter architectures and developmental gene regulation. Semin Cell Dev Biol 57, 11–23 (2016).

37. Ghandi, M., Lee, D., Mohammad-Noori, M. & Beer, M. A. Enhanced Regulatory Sequence Prediction Using Gapped k-mer Features. PLoS Comput Biol 10, e1003711 (2014).

38. Pampari, A. et al. ChromBPNet: bias factorized, base-resolution deep learning models of chromatin accessibility reveal cis-regulatory sequence syntax, transcription factor footprints and regulatory variants. bioRxiv 2024.12.25.630221 (2025) doi:10.1101/2024.12.25.630221.

39. Janssens, J. et al. Decoding gene regulation in the fly brain. Nature 601, 630–636 (2022).

40. De Winter, S., Konstantakos, V. & Aerts, S. Modelling and design of transcriptional enhancers. Nature Reviews Bioengineering (2025) doi:10.1038/s44222-025-00280-y.

41. Shrikumar, A., Prakash, E. & Kundaje, A. GkmExplain: Fast and accurate interpretation of nonlinear gapped k-mer SVMs. Bioinformatics 35, i173–i182 (2019).

42. Shrikumar, A. et al. Technical Note on Transcription Factor Motif Discovery from Importance Scores (TF-MoDISco) version 0.5.6.5. (2018).

43. Vierstra, J. et al. Global reference mapping of human transcription factor footprints. Nature 583, 729–736 (2020).

44. Spitz, F. & Furlong, E. E. M. Transcription factors: from enhancer binding to developmental control. Nat Rev Genet 13, 613–626 (2012).

45. Lambert, S. A. et al. Similarity regression predicts evolution of transcription factor sequence specificity. Nat Genet 51, 981–989 (2019).

46. Schep, A. N., Wu, B., Buenrostro, J. D. & Greenleaf, W. J. chromVAR: inferring transcription-factor-associated accessibility from single-cell epigenomic data. Nat Methods 14, 975–978 (2017).

47. Argelaguet, R. et al. Decoding gene regulation in the mouse embryo using single-cell multi-omics. bioRxiv 2022.06.15.496239 (2022).

48. Tournière, O. et al. NvPOU4/Brain3 Functions as a Terminal Selector Gene in the Nervous System of the Cnidarian Nematostella vectensis. Cell Rep 30, 4473–4489.e5 (2020).

49. Babonis, L. S. et al. Single-cell atavism reveals an ancient mechanism of cell type diversification in a sea anemone. Nat Commun 14, 885 (2023).

50. Steger, J. et al. Single-cell transcriptomics identifies conserved regulators of neuroglandular lineages. Cell Rep 40, 111370 (2022).

51. Jahnel, S. M., Walzl, M. & Technau, U. Development and epithelial organisation of muscle cells in the sea anemone Nematostella vectensis. Front Zool 11, 44 (2014).

52. Steinmetz, P. R. H., Aman, A., Kraus, J. E. M. & Technau, U. Gut-like ectodermal tissue in a sea anemone challenges germ layer homology. Nat Ecol Evol 1, 1535–1542 (2017).

53. Nakanishi, N., Renfer, E., Technau, U. & Rentzsch, F. Nervous systems of the sea anemone Nematostella vectensis are generated by ectoderm and endoderm and shaped by distinct mechanisms. Development 139, 347–357 (2012).

54. Lemaître, Q. I. B. et al. NvPrdm14d-expressing neural progenitor cells contribute to non-ectodermal neurogenesis in Nematostella vectensis. Nat Commun 14, 4854 (2023).

55. Cole, A. G. et al. Muscle cell-type diversification is driven by bHLH transcription factor expansion and extensive effector gene duplications. Nat Commun 14, 1747 (2023).

56. Hecker, N. et al. Enhancer-driven cell type comparison reveals similarities between the mammalian and bird pallium. Science *(1979)* 387, (2025).

## Methods references

57. Hand, C. & Uhlinger, K. R. The Culture, Sexual and Asexual Reproduction, and Growth of the Sea Anemone Nematostella vectensis. Biol Bull 182, 169–176 (1992).

58. Fritzenwanker, J. H. & Technau, U. Induction of gametogenesis in the basal cnidarian Nematostella vectensis(Anthozoa). Dev Genes Evol 212, 99–103 (2002).

59. Corces, M. R. et al. An improved ATAC-seq protocol reduces background and enables interrogation of frozen tissues. Nat Methods 14, 959 (2017).

60. De Rop, F. V et al. Hydrop enables droplet-based single-cell ATAC-seq and single-cell RNA-seq using dissolvable hydrogel beads. 11 (2022) doi:10.7554/eLife.

61. Drokhlyansky, E. et al. The Human and Mouse Enteric Nervous System at Single-Cell Resolution. Cell 182, 1606–1622.e23 (2020).

62. Torres-Méndez, A. et al. A novel protein domain in an ancestral splicing factor drove the evolution of neural microexons. Nat Ecol Evol 3, 691–701 (2019).

63. Najle, S. R. et al. Stepwise emergence of the neuronal gene expression program in early animal evolution. Cell 186, 4676–4693.e29 (2023).

64. García-Castro, H. et al. ACME dissociation: a versatile cell fixation-dissociation method for single-cell transcriptomics. Genome Biol 22, 89 (2021).

65. Yu, W., Uzun, Y., Zhu, Q., Chen, C. & Tan, K. scATAC-pro: a comprehensive workbench for single-cell chromatin accessibility sequencing data. Genome Biol 21, 94 (2020).

66. Fletcher, C. & Pereira da Conceicoa, L. The genome sequence of the starlet sea anemone, Nematostella vectensis (Stephenson, 1935). Wellcome Open Res 8, 79 (2023).

67. Li, H. & Durbin, R. Fast and accurate short read alignment with Burrows–Wheeler transform. Bioinformatics 25, 1754–1760 (2009).

68. Lun, A. T. L. et al. EmptyDrops: distinguishing cells from empty droplets in droplet-based single-cell RNA sequencing data. Genome Biol 20, 63 (2019).

69. Granja, J. M. et al. ArchR is a scalable software package for integrative single-cell chromatin accessibility analysis. Nat Genet 53, 935–935 (2021).

70. Persad, S. et al. SEACells infers transcriptional and epigenomic cellular states from single-cell genomics data. Nat Biotechnol 2022.04.02.486748 (2023) doi:10.1038/s41587-023-01716-9.

71. van den Oord, J., et al. SCENIC: single-cell regulatory network inference and clustering. Nat Methods 14, 1083–1086 (2017).

72. Zhang, Y. et al. Model-based analysis of ChIP-Seq (MACS). Genome Biol 9, R137 (2008).

73. Huber, B. R. & Bulyk, M. L. Meta-analysis discovery of tissue-specific DNA sequence motifs from mammalian gene expression data. BMC Bioinformatics 7, 1–25 (2006).

74. Schep, A. N., Wu, B., Buenrostro, J. D. & Greenleaf, W. J. chromVAR: inferring transcription-factor-associated accessibility from single-cell epigenomic data. Nat Methods 14, 975–978 (2017).

75. Argelaguet, R. et al. Decoding gene regulation in the mouse embryo using single-cell multi-omics. bioRxiv 2022.06.15.496239 (2022).

76. Machlab, D. et al. monaLisa: an R/Bioconductor package for identifying regulatory motifs. Bioinformatics 1–2 (2022) doi:10.1093/bioinformatics/btac102.

77. Lee, D. LS-GKM: a new gkm-SVM for large-scale datasets. Bioinformatics 32, 2196–2198 (2016).

78. Pampari, A. et al. ChromBPNet: bias factorized, base-resolution deep learning models of chromatin accessibility reveal cis-regulatory sequence syntax, transcription factor footprints and regulatory variants. bioRxiv 2024.12.25.630221 (2025) doi:10.1101/2024.12.25.630221.

79. Shrikumar, A., Greenside, P. & Kundaje, A. Learning Important Features Through Propagating Activation Differences. (2017).

80. Shrikumar, A. et al. Technical Note on Transcription Factor Motif Discovery from Importance Scores (TF-MoDISco) version 0.5.6.5. (2018).

81. Renfer, E. & Technau, U. Meganuclease-assisted generation of stable transgenics in the sea anemone Nematostella vectensis. Nat Protoc 12, 1844–1854 (2017).

